# Exploring the Impact of Analysis Software on Task fMRI Results

**DOI:** 10.1101/285585

**Authors:** Alexander Bowring, Camille Maumet, Thomas E. Nichols

**Author notes:** These senior authors contributed equally to this study (Camille Maumet, Thomas E. Nichols). Address for correspondence: T. Nichols, Big Data Institute Building, Old Road Campus, Roosevelt Drive, Oxford OX3 7LF UK.

## Abstract

A wealth of analysis tools are available to fMRI researchers in order to extract patterns of task variation and, ultimately, understand cognitive function. However, this ‘methodological plurality’ comes with a drawback. While conceptually similar, two different analysis pipelines applied on the same dataset may not produce the same scientific results. Differences in methods, implementations across software packages, and even operating systems or software versions all contribute to this variability. Consequently, attention in the field has recently been directed to reproducibility and data sharing. Neuroimaging is currently experiencing a surge in initiatives to improve research practices and ensure that all conclusions inferred from an fMRI study are replicable.

In this work, our goal is to understand how choice of software package impacts on analysis results. We use publicly shared data from three published task fMRI neuroimaging studies, reanalyzing each study using the three main neuroimaging software packages, AFNI, FSL and SPM, using parametric and nonparametric inference. We obtain all information on how to process, analyze, and model each dataset from the publications. We make quantitative and qualitative comparisons between our replications to gauge the scale of variability in our results and assess the fundamental differences between each software package. While qualitatively we find broad similarities between packages, we also discover marked differences, such as Dice similarity coefficients ranging from 0.000 - 0.684 in comparisons of thresholded statistic maps between software. We discuss the challenges involved in trying to reanalyse the published studies, and highlight our own efforts to make this research reproducible.

## 1. Introduction

Functional Magnetic Resonance Imaging (fMRI) for human brain mapping gives researchers remarkable power to probe the underpinnings of human cognition, behavior and emotion. As an active field of research for over 25 years, there are now a multitude of ways to analyze a single neuroimaging study. The plethora of techniques and tools available are a platform from which we have the potential to gain remarkable insight into how the human brain works. However, high analytic flexibility has also been pinpointed as a key factor that can lead to false-positive and non-reproducible results (Ioannidis, 2005; Wager et al., 2009). Because of this, neuroimagers must be particularly judicious: choice of analysis pipeline, operating system and even software version may influence the final research outcome of a study.

The extent to which varying processing conditions can lead to discrepancies in observed results has been highlighted throughout the neuroimaging literature. In a study examining the use of FreeSurfer to measure the cortical thickness and volume of structural brain images (Gronenschild et al., 2012), a change in software version was shown to lead to increases of over 10% in observed anatomical measurement; a switch in workstation from which the software was run also manifested significant deviations in the final result. In related work (Glatard et al., 2015), changes in operating system lead to differences in the results of an independent component analysis of resting state fMRI data carried out using FSL. Here, disparities in both the number of components determined as well as information between matched components were found when the analysis was conducted on two separate computing clusters. For task-based fMRI, the impact of methodological choices has been investigated extensively. Choices for each individual procedure in the analysis pipeline (for example, head-motion regression (Lund et al., 2005), temporal filtering (Skudlarski et al., 1999), and autocorrelation correction (Woolrich et al., 2001)) alongside the order in which these procedures are conducted (Carp, 2013) can deeply influence the final determined areas of brain activation. In perhaps the most comprehensive of such studies (Carp, 2012a), a single publicly available fMRI dataset was analyzed using over 6,000 unique analysis pipelines, generating 34,560 unique thresholded activation images. These results displayed a substantial degree of flexibility in both the sizes and locations of significant activation. In combination, these examples of research shape a sombre picture for the possibility of study reproducibility.

While each of the aforementioned studies investigated the effect of either software version, operating system, or analysis pipeline on analytic variability, the choice of software package for carrying out the analysis remained fixed in each study. This is despite a vast array of analysis packages that are now freely available to researchers. The three most popular of these packages for fMRI data analysis are AFNI (RRID:SCR_005927; (Cox, 1996)), FSL (RRID:SCR_002823; (Jenkinson et al., 2012)) and SPM (RRID:SCR_007037; (Penny et al., 2011)). While SPM is the oldest, FSL has grown in popularity and together the three packages have been estimated to account for 80% of published functional neuroimaging results (Carp, 2012b). Although there are differences in how each software package models and processes data, the analysis framework for task fMRI - now a mature research area - is expected to be similar across software, and hence the results yielded from each package should be comparable. We therefore seek to answer the question: How much of the variability in neuroimaging results is attributable to the choice of analysis software package?

In this work we reanalyze data from three published neuroimaging studies using each of the three main software packages and quantify differences in the results. We choose three publications with data that have been made publicly available on the OpenfMRI database (RRID:SCR_005031, http://openfmri.org; (Poldrack et al., 2013)), recently relaunched as OpenNeuro (http://openneuro.org), and attempt to recreate the main figure from each publication by replicating the original analysis within each package. These particular studies were selected on the basis that they reported clearly defined regions of brain activation and utilized analysis procedures feasible across the three software packages. We then make a number of comparisons to assess the similarity of our results. While a similar study from our group has explored the results produced by each of these packages after implementing analysis pipelines using the default settings in each software (Pauli et al., 2016), here we attempt to make the analysis pipelines as similar as possible while still maintaining comparability across the three packages. While our primary focus is comparing standard results across software, we also aim to address recent concerns about the multiple-testing-corrected parametric inferences that each of these studies used (Eklund et al., 2016). For each study, we also conduct equivalent inference procedures (when possible) using nonparametric statistics in each package.

Although our work has been primarily designed to understand the differences between software packages, we also see this as an exercise in computational reproducibility (Peng, 2011). In recent years, a number of initiatives and guidelines (Poldrack et al., 2017) have materialized to ensure research is conducted in an open and transparent fashion. For each of our analyses, we confine ourselves to the respective publication for all information on how to process and model the data. We discuss the challenges involved in this process, and evaluate whether our reanalyses are a success by comparing our results to those given in the main figure of the respective publication. Great care has also been taken to ensure all figures and results presented here are themselves reproducible; we describe the scripts, notebooks and other tools used to make this possible which we believe are highly generalizable across neuroimaging studies.

## 2. Methods

### 2.1 Study Description and Data Source

We selected three functional fMRI studies for reanalysis from the publicly accessible OpenfMRI data repository: ds000001 (Revision: 2.0.4; (Schonberg et al., 2012)), ds000109 (Revision 2.0.2; (Moran et al., 2012)), and ds000120 (Revision 1.0.0; (Padmanabhan et al., 2011)). Each of the datasets have been organized in compliance with the Brain Imaging Data Structure (BIDS, RRID:SCR_016124; (Gorgolewski et al., 2016)). These datasets were chosen following an extensive selection procedure (carried out between May 2016–November 2016), whereby we vetted the associated publication for each dataset stored in the repository. We sought studies with simple analysis pipelines and clearly reported regions of brain activation that would be easily comparable to our own results. Exclusion criteria included the use of custom software, activations defined using small volume correction, and application of more intricate methods such as region of interest and robust regression analysis, which we believed could be impractical to implement across all analysis software. A full description of the paradigm for each of our chosen studies is included in the respective publication; here we give a brief overview.

For the ds000001 study, 16 healthy adult subjects participated in a balloon analog risk task over three scanning sessions. On each trial, subjects were presented with a simulated balloon, and offered a monetary reward to ‘pump’ the balloon. With each successive pump the money would accumulate, and at each stage of the trial subjects had a choice of whether they wished to pump again or cash-out. After a certain number of pumps, which varied between trials, the balloon exploded. If subjects had cashed-out before this point they were rewarded with all the money they had earned during the trial, however if the balloon exploded all money accumulated was lost. Three different colored ‘reward’ balloons were used between trials, each having a different explosion probability, as well as a gray ‘control’ balloon, which had no monetary value and would disappear from the screen after a predetermined number of pumps. Here we reproduce the result contrasting the parametrically modulated activations of pumps of the reward balloons versus pumps of the control balloon, corresponding to Figure 3 and Table 2 in the original paper.

The ds000109 study investigated the ability of people from different age-groups to understand the mental state of others. A total of 48 subjects were scanned, although 43 had acceptable data for the false belief task - 29 younger adults and 14 older adults. In this task participants listened to either a ‘false belief’ or ‘false photo’ story. A false belief story would entail an object being moved from one place to another, with certain characters witnessing the change in location while others were unaware. False photo stories were similar except involved some physical representation, such as a photo of an object in a location from which it had been subsequently removed. The task had a block design where stories were represented for ten seconds, after which participants had to answer a question about one of the character’s perceptions of the location of the object. We reproduce the contrast map of false belief versus false photo activations for the young adults, corresponding to Figure 5a and Table 3 from the original publication.

Finally, the the ds000120 study explored reward processing across different age groups. fMRI results are reported on 30 subjects, with 10 participants belonging to each of the three age groups (children, adolescents and adults). Participants took part in an antisaccade task where a visual stimuli was presented in each trial and subjects were instructed to quickly fixate their gaze on the side of the screen opposite to the stimuli. Prior to a trial, subjects were given a visual cue to signal whether or not they had the potential to win a monetary reward based on their upcoming performance (a ‘reward’ or ‘neutral’ trial). In this paper we reproduce the main effect of time activation map - an F-statistic for any non-zero coefficients in the sine HRF basis - corresponding to Figure 3 and Table 1 in the original publication.

**Table 1.**
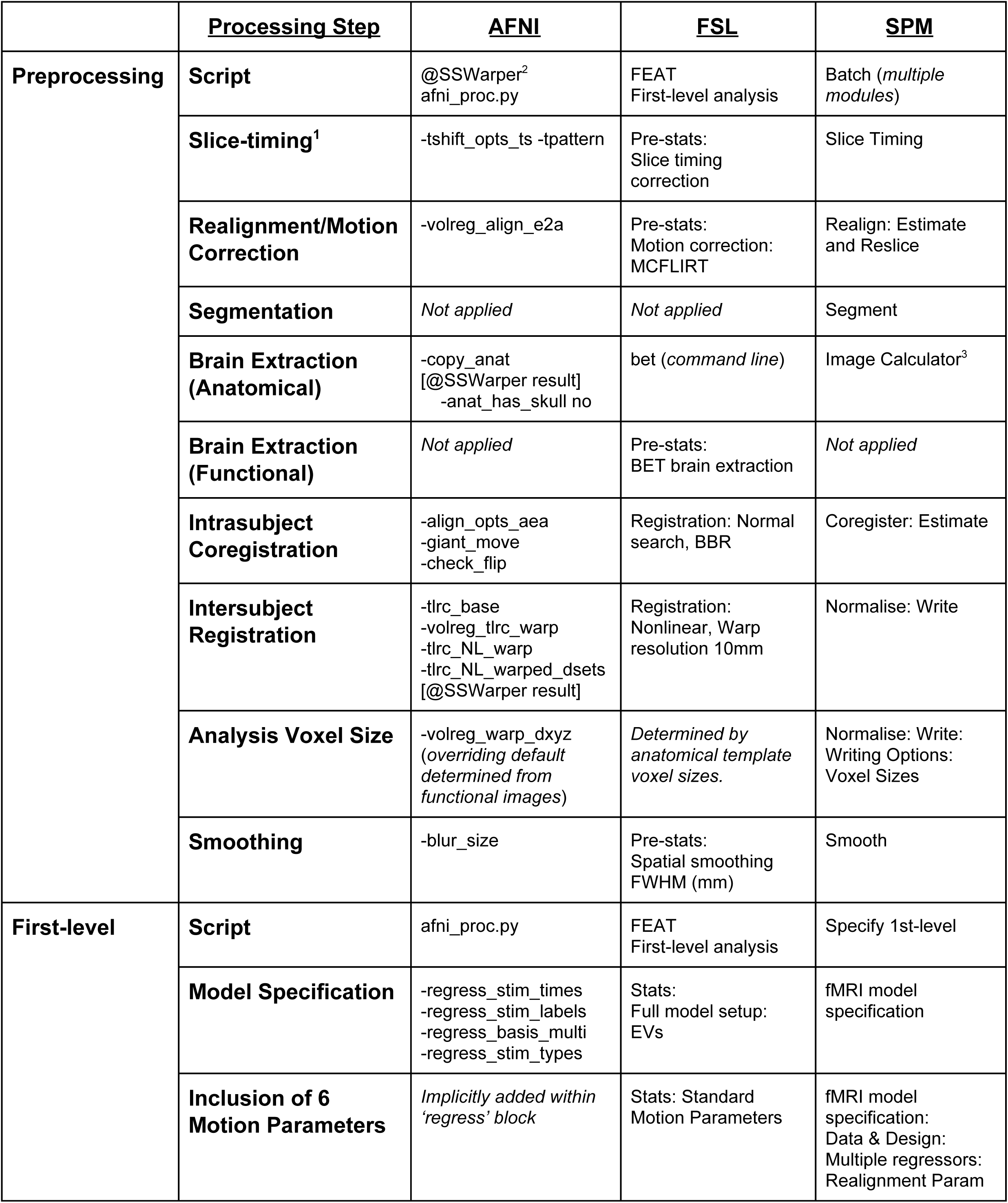
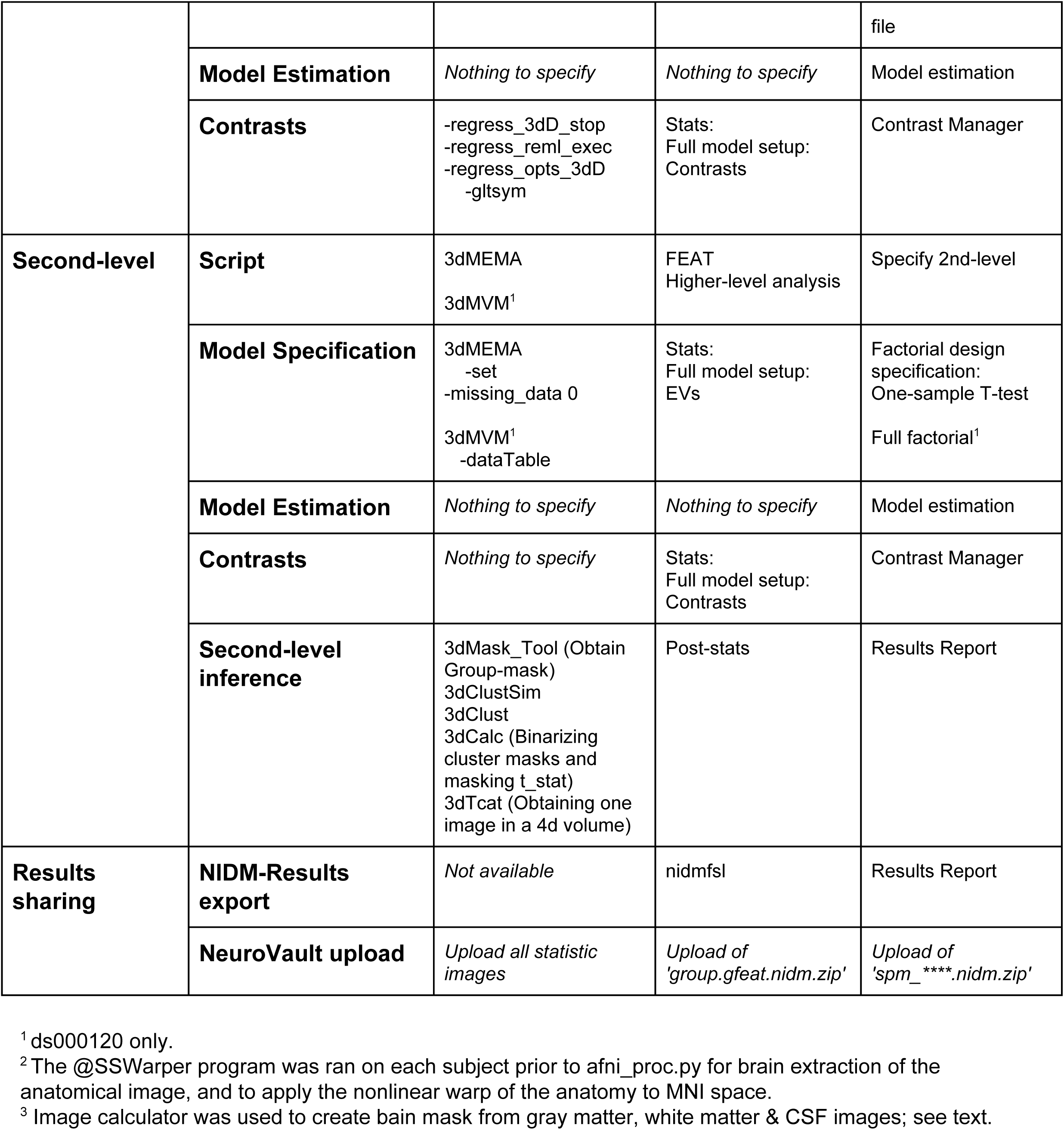
Software Processing Steps. Implementation of each of the processing steps (ds000001, ds000109, ds000120) within AFNI, FSL and SPM.

### 2.2 Data Analyses

All data analyses were conducted using AFNI (version AFNI_17.0.18), FSL (version 5.0.10), and SPM (version SPM12, v6906). Computation was performed on a cluster comprised of 12 Dell PowerEdge servers (6 R410, 12 core 2.40GHz processors, 6 R420, 12 core 2.80GHz processors) running CentOS 7.3.

#### 2.2.1 Pipeline

A full decomposition of the pipelines implemented within the three packages for each study is presented in Table 1. Here, we give a brief description of the procedures.

In AFNI, preprocessing and subject-level analyses were conducted using the @SSwarper program and afni_proc.py. For ds000001 and ds000109, we used the 3dMEMA program to perform a one-sample T-test, while for ds000120 we used the 3dMVM program at the second level to conduct a mixed-effects analysis, generating an F-statistic for the main effect of time.

In FSL, analyses were carried out using the FMRI Expert Analysis Tool (FEAT, v6.00). For each analysis, at the first level a separate .fsf file was created for each scanning session. Runs were then combined as part of a second level fixed-effects model, yielding results which were subsequently inputted into a group analysis.

In SPM, preprocessing, subject-and group-level analyses were conducted by selecting the relevant modules within SPM’s Batch Editor. In particular, subject-level and group-level analyses were conducted using the Specify 1st-level and Specify 2nd-level modules respectively.

Once analyses were complete, the results for each software package were exported as NIDM-Results packs (FSL and SPM only, (Maumet et al., 2016)) and uploaded to a public collection on the NeuroVault (RRID:SCR_003806, http://neurovault.org; (Gorgolewski et al., 2015)) online data repository.

#### 2.2.2 Common Processing Steps

A number of processing steps for each package were included in all of our analyses, regardless of whether they had been implemented in the original study. While this meant deviating from an exact replication of the original pipeline, these processing steps were either fundamental to ensure that results from each software package could be compared objectively, or steps that are widely accepted as best practice within the community. In this section we describe these steps.

Successful coregistration of the functional data to the structural brain images - and subsequently - registration to the MNI template, was of paramount importance to us for fair comparability of the results. During our first attempt at analysing the ds000001 dataset we discovered that seven subjects had essential orientation information missing from the NIfTI header fields of their functional and structural data. As the source DICOM files were no longer available, the original position matrices for this dataset were unable to be retrieved. This caused coregistration to fail for several subjects across all three software packages in our initial analysis of this data. We rectified the issue by manually setting the origins of the functional and structural data. OpenfMRI released a revision (Revision: 2.0.4) of our amended dataset which we used for the analysis. Further to this, we also set a number of common preprocessing steps within each package to be applied in all our analyses.

Firstly, brain extraction was conducted on the structural image in all software. We did this to improve registration and segmentation. In AFNI, brain extraction was carried out using 3dskullstrip, that was called implicitly from within the @SSwarper program. The skull-stripped anatomical volume obtained here was inputted into our afni_proc.py scripts where further preprocessing and first-level analyses were carried out. In FSL, brain extraction was performed on both the functional and structural data. The Brain Extraction Tool (BET; (Smith, 2002)) was applied to each structural image from the command line before preprocessing, and for functional data with the BET option within the Pre-stats module of FEAT. In SPM, brain extraction was implemented via the segmented structural images. Gray matter, white matter and CSF images were summed and binarised at 0.5 to create a brain mask, which was applied to the bias corrected structural image using the Image Calculator.

Coregistration of the functional data to the anatomy was carried out for the most part using the default settings in each software. In AFNI, alignment of the data was conducted using the align_epi_anat.py program called implicitly from the align block within the afni_proc.py scripts. We included the -volreg_align_e2a option within our scripts to specify alignment of the functional data onto the anatomy, as by default AFNI conducts the inverse transformation of anatomy onto functional. Further to this, we also added the -align_opts_aea program to all of our scripts with the -giant_move and -check_flip options to allow for larger transformations between the images. In FSL, coregistration was carried out within FEAT using the default linear registration methods with a Boundary-Based Registration (BBR) cost function. The default methods were also applied within SPM’s Coregister: Estimate module, using a normalised mutual information cost function.

Registration of the structural and functional data to the anatomical template was executed using each packages nonlinear settings. In AFNI, nonlinear registration of the anatomical data to the MNI template was conducted as part of the @SSwarper program ran prior to the afni_proc.py script. The warps computed by @SSwarper were passed to afni_proc.py using the -tlrc_NL_warpred_dsets option, and applied to the functional data within the tlrc block using the -volreg_tlrc_warp option. By default, the resampled functional data in MNI space has voxel size determined from the raw 4D data; we forced 2mm cubic voxels with the -volreg_warp_dxyz option for compatibility with FSL and SPM’s 2mm default. In FSL, registration to the MNI template was conducted using FMRIB’s Nonlinear Image Registration Tool (FNIRT; (Andersson et al., 2007)), controlling the degrees of freedom of the transformation with a warp resolution of 10mm. In SPM, the nonlinear deformations to MNI space were obtained as part of the Segment module and then applied to the structural and functional data within the Normalise: Write module.

As a form of quality control, we created mean and standard deviation images of the subject-level MNI-transformed anatomical and mean functional images. Alongside the subject-level data, these images were assessed to check that registration to MNI space had been successful. When intersubject registration failed remedial steps were taken within each software; these are described in the software implementation parts of the following study-specific analysis sections.

Finally, across all software packages six motion regressors were included in the analysis design matrix to regress out motion-related fluctuations in the BOLD signal. Use of six or more derived motion regressors is commonly recommended as good practice, and we chose to use just six regressors as this could be easily implemented across software.

We now describe the task-specific analysis procedures for each of the three studies as carried out in the original publications, and how these methods were implemented within each package. While we decided to keep the above steps of the analysis pipelines fixed, for all remaining procedures we attempted to remain true to the original study. Any further deviations necessitated are discussed in the software implementation sections. Notably, apart from the addition of six motion regressors, *all* of our common steps relate to preprocessing, and hence for first-and group-level analysis we attempt to exactly replicate the original study.

#### 2.2.3 ds000001 Analyses

In the publication associated with the ds000001 study all preprocessing and analysis was conducted within FSL (version 4.1.6). Data on all 16 subjects were available to us on OpenfMRI. In the original preprocessing, the first two volumes of the functional data were discarded and the highpass-filter was set to a sigma of 50.0s. Motion correction was conducted using MCFLIRT and brain extraction of the functional data was applied with BET, after which FSL’s standard three-step registration procedure was carried out to align the functional images to the structural scan. Spatial normalization was implemented with FMRIB’s Linear Image Registration Tool (FLIRT; (Jenkinson et al., 2002)), and data were smoothed using a 5mm full-width-half-maximum (FWHM) Gaussian kernel. At the run level, each of the events were convolved using a canonical double-gamma haemodynamic response function (HRF); FEAT’s (then newly available) outlier de-weighting was used. Subject-level analysis of the functional data were conducted using a general linear model (GLM) within FEAT, where a selection of the regressors were orthogonalized. The three scanning sessions for each participant were carried out separately and then combined together at the second level. A pair of one-sided T-tests were conducted at the group-level to test for positive and negative effects separately. For each test, clusterwise inference was performed using an uncorrected cluster-forming threshold of p < 0.01, FWE-corrected clusterwise threshold of p < 0.05 using Gaussian random field theory.

We opted to not use outlier de-weighting on the basis that such methods were impractical to implement across all software packages.

#### AFNI Implementation

Using our default procedure for the AFNI analysis, we found that coregistration of the functional scans onto the anatomy failed for four subjects. To remedy this issue, for this study we modified our afni_proc.py scripts: Within the -align_opts_aea module, the ‘-ginormous move’ option was added to align centers of the functional and anatomical volumes, and the ‘-cost lpc+ZZ’ option was used to apply a weighted combination of cost functionals. Both of these changes are recommended for data with little structural detail. Following these modifications all coregistrations were successful.

To replicate the orthogonalization methods from the original study, a separate orthogonalization script was ran for each subject prior to the first-level analyses. Within this script, the (un-orthogonalized) regressors were generated by passing the event timing files to 3dDeconvolve, after which the 3dTproject command was used to obtain the desired projections. The orthogonalized regressor files outputted from this script were then entered into afni_proc.py to replicate the original subject-level analysis model.

Trials were convolved with a single gamma HRF using either the BLOCK or dmBLOCK option within the-regress_basis_multi module, determined by whether the event file had fixed or variable duration times respectively. The -regress_stim_types option was added to our afni_proc.py script to specify event files for regressors which had been parametrically modulated in the original study, and identify the orthogonalized regressors.

At the group level, we performed a mixed-effects analysis using 3dMEMA. The critical cluster size threshold was determined by Monte Carlo simulation with the 3dClustSim program.

#### FSL Implementation

Implementation in FSL closely followed the original procedure described above, with the exception that nonlinear registration was used to transform the data to standard space.

#### SPM Implementation

Implementation in SPM closely followed the pipeline outlined in Table 1.

#### 2.2.4 ds000109 Analyses

The original preprocessing and statistical analysis for the ds000109 study was carried out using SPM8. Data were shared on 36 of the 40 subjects, 21 of which were young adult subjects that had fMRI data compatible for our reanalysis. First, functional data were realigned and unwarped to correct for head motion and geometric distortions. After transforming the data into a standardized space, the normalized data were smoothed with an 8mm FWHM Gaussian kernel. Further to this, custom software was applied to exclude functional volumes where head motion had exceeded a certain limit, however this process was omitted from our pipelines since this feature was not available in any of the software packages. The preprocessed data were entered into a GLM for first level analysis where trials were modeled using a block design and convolved using SPM’s canonical HRF. Each participant’s contrast images were then entered into a one-sample group analysis using clusterwise inference, cluster forming threshold of p < 0.005, 5% level FWE using random field theory; in their analysis, this amounted to a critical cluster size threshold of 56 voxels.

#### AFNI Implementation

Intersubject registration to the MNI atlas failed for one subject, for which part of the frontal lobe was missing. We addressed this by revising this study’s AFNI pipeline to use the -pad_base 60 option within the -tlrc_opts_at module included in afni_proc.py. This gave extra padding to the MNI template so that no part of the functional image was lost during the alignment.

The HRF was modelled with SPM’s canonical HRF using the SPMG1 option for each event within the -regress_basis_multi option and passing the duration of the regressor as an argument to the function.

At the group level, we performed a mixed-effects analysis using 3dMEMA. P-values were determined by Monte Carlo simulations with 3dClustSim.

#### FSL Implementation

To recreate the original HRF model in FSL, we chose the Double-Gamma HRF from the convolution options within FEAT.

#### SPM Implementation

Implementation in SPM closely followed the original procedure described above.

#### 2.2.5 ds000120 Analyses

A multi-software analysis procedure was used for the ds000120 study, where data were preprocessed with FSL and then analyzed using AFNI. fMRI data were shared on OpenfMRI for 26 of the original 30 subjects, and 17 had data available on the task of interest. This was the only study that applied slice-timing correction, adjusting the functional data for an interleaved slice acquisition. Functional scans were realigned to the middle volume, and following brain extraction with BET, registered to the structural scan in Talairach space using FLIRT and FNIRT. Data were high-pass filtered with a sigma value of 30.0s and smoothed with a 5mm FWHM Gaussian kernel. Like the previous study, further methods were used to remove functional volumes with excessive motion which have been left out from our analyses due to discordance across software. Subject-level analysis was conducted within AFNI. To allow for flexible modelling of the response to the saccade task, this study used a HRF basis consisting of eight sine functions with a post-stimulus window length of 24.0s. At the group level, subjects were entered into a mixed-effect model, with subjects as a random factor, trial type (reward, neutral) and time as within-group factors, and age group (child, adolescent, adult) as a between-group factor. Clusterwise inference was used on the main effect of time activation map (F_{8,142} statistic), cluster-forming threshold of p < 0.001, controlling FWE at the 5% level, obtained with Monte Carlo methods. This computed critical cluster size threshold was 23 voxels.

#### AFNI Implementation

Slice timing was conducted using the -tshift_opts_ts program within afni_proc.py with the -tpattern option applied to specify an interleaved slice acquisition.

The sine basis set used for the HRF was modelled using the -regress_basis_multi module with the SIN option.

At the group level, a mixed-effect analysis was carried out with the 3dMVM program. Following this, 3dClustSim was used to obtain the cluster extent corresponding to the original study threshold. In our analysis we found the cluster size threshold to be 48 voxels.

#### FSL Implementation

The repeated-measures design used in the group-level analysis of the original study was not feasible to implement in FSL, and as such, we did not attempt an FSL reanalysis for this study. (The FEAT manual does describes “Repeated Measures” examples, but these are based on a restrictive assumption of compound symmetry; here this would entail assuming that all 8*7/2=28 correlations among the basis regression coefficients are equal.)

#### SPM Implementation

Slice timing was conducted using the Slice Timing module within the Batch Editor of SPM.

Although an exact equivalent of the original HRF model was not possible in SPM, we chose the closest equivalent using the Fourier basis set with an order of 4, leading to a total of 9 basis functions fit to each of the reward & neutral conditions for each of the three runs. A set of 9 first level contrasts computed the average Fourier coefficients over conditions and runs.

To reproduce the group-level analysis in SPM, a full factorial design was chosen within the ‘Factorial design specification’ module of the Batch Editor, with a time factor (9 levels) and adding age-group to the model using two covariates (adolescent vs child, adult vs child); the main effect of time was tested with an F-contrast.

### 2.3 Comparison Methods

We applied three separate quantitative methods to measure the similarity between the group results obtained within each software package for each of the three studies.

Firstly, Bland-Altman plots comparing the unthresholded group statistic maps were created for each pairwise combination of software packages. These plotted the difference between the statistic values (y-axis) against the mean statistic value (x-axis) for all voxels lying inside the intersection of the two software’s analysis masks. In addition to this, we also created Bland-Altman plots to compare percentage BOLD change maps (for ds000120, partial R^2^ maps) between software. For each package, an appropriate normalization of the group-level beta maps was conducted to convert to percentage BOLD change units. Due to differences in how each package scales the data, a different normalization was required for each of the three packages. For ds000120, the partial R^2^ maps were computed via a transformation of the group-level F-statistic images. We provide full details on how each of these procedures were carried out in the appendix (Appendix 1. for percent BOLD change, Appendix 2. for partial R^2^).

We also computed the Dice similarity coefficient for each pairwise combination of the group-level thresholded statistic maps. The coefficient is calculated as the cardinality of the intersection of the thresholded maps divided by the average of the cardinality of each thresholded map. While Bland-Altman is interested in the similarity between statistic values, Dice measures the overlap of voxels as a means to assess the *spatial similarity* of activated clusters. The coefficient takes a value between zero and one, where one indicates complete congruence between the size and location of clusters in both thresholded maps, while zero indicates no agreement. Dice coefficients were computed over the intersection of the pair of analysis masks, to assess only regions where activation could occur in both packages. We also calculated the percentage of ‘spill over’ activation, i.e. the percentage of activation in one software’s thresholded statistic map that fell outside of the analysis mask of the other software.

A particular concern we had was that a pair of statistic images could in essence be very similar, but differ by a scale factor over all voxels. Another possibility was that one software could have greater sensitivity for voxels where signal was present, causing differences between images only for relatively higher statistical values. Both of these features would not be identifiable using our previous comparison methods. To address this, we computed the Euler Characteristic (EC) for each software’s group T-statistic map (F-statistic for ds000120), thresholded using T-values between -6 to 6 (0 to 6 for ds000120; increasing with an increment of 0.2). Alongside the EC, we also computed the number of clusters in the statistic images using the same thresholds. For a given threshold t, the EC calculates the number of clusters minus the numbers of ‘handles’ plus the number of ‘holes’ in the thresholded image. For large t, we expect the handles and holes to disappear, and therefore the EC provides an approximation of the number of clusters in an image. For smaller t, we expect our thresholded image to be one connected cluster with many holes and handles (like Swiss cheese) - it is in this situation where the EC is clearly more informative about differences between images than the cluster count alone. Over all t, the EC curve provides a signature of an entire statistic image, and provides a means to assess whether only superficial scaling differences are responsible for disparities between a pair of images.

For a qualitative assessment of whether similar activation patterns were displayed between packages, a NeuroSynth (RRID:SCR_006798, http://neurosynth.org) association analysis was conducted on each software’s unthresholded statistic maps. These analyses performed a cognitive decoding of the unthresholded statistic image with images in the NeuroSynth database, to find the words or phrases most strongly associated with the activation patterns found in the statistic map.

Finally, we visually compared the corresponding slices of each software’s thresholded statistic map to those presented in the publication figure we had attempted to recreate. Ensuring we had found activation in approximately the same regions as the original publication gave us an indication that we had successfully replicated the study’s analysis pipeline.

### 2.4 Permutation Test Methods

For ds000001 and ds000109, in parallel to our replication analyses we computed an additional set of group-level results applying nonparametric permutation test inference procedures available within each software package (a one-sample repeated measures permutation test needed for ds000120 was not available in AFNI). The first level contrast maps obtained from our initial replications for each subject were entered into a group-level one-sample T-test where clusterwise inference was conducted using the same cluster-forming thresholds, and then 5% level FWE corrected thresholds were computed by permutation, using 10,000 permutations.

#### AFNI Implementation

In AFNI, permutation inference was carried out using the 3dttest++ module with the -ClustSim option. By applying this option, permutation generated noise realisations which 3dClustSim used to generate cluster-threshold tables. Significant clusters in the group-activation map were found with 3dclust, using a critical cluster size threshold extracted from the 3dClustSim output.

#### FSL Implementation

Permutation test inference was conducted in FSL using randomise version 2.9 (Winkler et al., 2014). This outputted a ‘corrp’ image which was then used to mask the raw T-statistic image to show significant voxels for the appropriate thresholds.

#### SPM Implementation

The Statistical nonParametric Mapping (SnPM, version SnPM13; RRID:SCR_002092; (Nichols and Holmes, 2002)) toolbox was used to carry out permutation tests in SPM. The “MultiSub: One Sample T test on diffs/contrasts”, Compute and Inference modules within SnPM were applied to obtain the final group-level activation maps.

Each of the comparison methods described in the previous section were also applied to our permutation results to assess cross-software differences for nonparamteric inference methods. In addition, we also generated intra-software Bland-Altman plots and Dice coefficients to understand differences between the parametric and nonparametric methods applied within each package. These methods were excluded for ds000120, since it was not possible to conduct permutation inference for an F-test within AFNI.

### 2.5 Scripting of analyses and figures

AFNI and FSL scripts were written in Python 2.7.14 and SPM scripts were written in Matlab R2016b. Scripts were made generalizable, such that the only study-specific differences for each of the analyses in a software package were the raw data and working directory inputs, subject- and group-level analysis templates (as well as a run-level template for FSL), and a unique conditions structure necessary for creating the onset files for the specified study. For each analysis package, a script was written to extract the stimulus timings from the raw data to create event files that were compatible within the software. Subject-level analysis templates were batch scripts created for each study containing all processing steps of the subject analysis pipeline for the respective software, with holding variables used where subject-or run-specific inputs were required. The main script would take the template as an input, and cycling through each of the subjects, replace the holding variables with appropriate pathnames to create distinct batch scripts for each subject. These were then executed to obtain subject-level results for all participants in the study.

A Python Jupyter Notebook (Kluyver et al., 2016) was created for each of the three studies. Each notebook harvests our results data from NeuroVault and applies the variety of methods discussed in the previous section using NiBabel 2.2.0 (Brett et al., 2017), NumPy 1.13.3 (Walt et al., 2011) and Pandas 0.20.3 (McKinney and Others, 2010) packages. Figures were created using Matplotlib 2.1.0 (Hunter, 2007) and Nilearn 0.4.0 (Abraham et al., 2014).

## 3. Results

All scripts and results are available through our Open Science Framework (OSF)(Erin D. Foster, 2017) Project at https://osf.io/U2Q4Y/ (Alex Bowring et al., 2018), and group-level statistic maps used to create the figures in this section are available on NeuroVault: https://neurovault.org/collections/4110/, https://neurovault.org/collections/4099/, https://neurovault.org/collections/4100/, for ds0000001, ds000109 and ds000120 respectively. All analysis scripts, results reports, and notebooks for each study are available through Zenodo (Nielsen and Smith, 2014) at https://doi.org/10.5281/zenodo.1413540 (Alexander Bowring et al., 2018, Software_Comparsion: Version 0.4).

Registration of each subject’s functional data onto the anatomy was visually assessed. Mean and standard deviation images of the structural and (mean) functional data (Fig. S1) are remarkably similar.

### 3.1 Cross-Software Variability

While qualitatively similar, variability in T-statistic values and locations of significant activation was substantial between software packages across all three studies.

Comparisons of the thresholded results with the published findings are shown in Figure 1a, with further multi-slice comparisons across software in Figure 1b (also in Figs. S2, S4, & S6). The ds000001 study described positive activation in the bilateral anterior insula, dorsal anterior cingulate cortex (ACC), and right dorsolateral prefrontal cortex, and negative activation in the ventromedial prefrontal cortex and bilateral medial temporal lobe. In our reanalysis (Fig. 1a, left) all three software found activation in these set of regions, with the exception that decreases in the medial temporal lobe were unilateral in FSL and SPM (left only). FSL also detected a visual response that neither AFNI or SPM picked up on (Fig. 1a, left, and Fig. 1b, top, z = 0 slice).

**Figure 1a.**
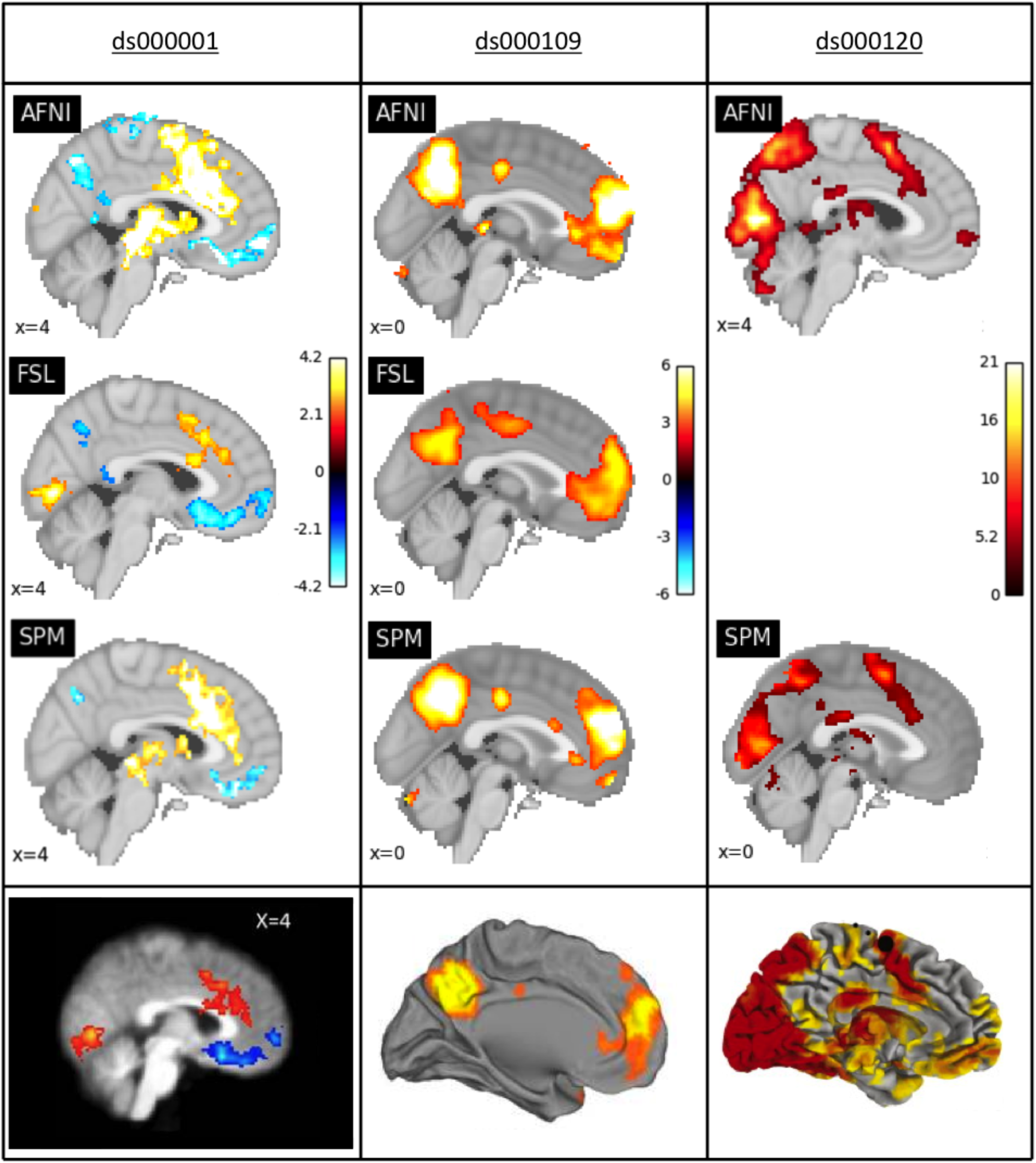
Comparison of the thresholded statistic maps from our reanalysis with the main figures from each of the three publications. Left: For ds000001 data, thresholded T-statistic images contrasting the parametric modulation of pumps of reward balloons versus the parametric modulation of the control balloon; beneath, a sagittal slice taken from Figure 3 in Schonberg et al (2012). Middle: For ds000109, thresholded T-statistic maps of the false belief vs false photo contrast; beneath, a midsagittal render from Moran et al (2012). Right: For ds000120, thresholded F-statistic images of the main effect of time contrast; beneath, a midsagittal render from Figure 3 in Padmanabhan et al. (2011). Note that for ds000109 and ds000120 the publication’s figures are renderings onto the cortical surface while our results are slice views. While each major activation area found in the original study exists in the reanalyses, there is substantial variation between each reanalysis.

**Figure 1b.**
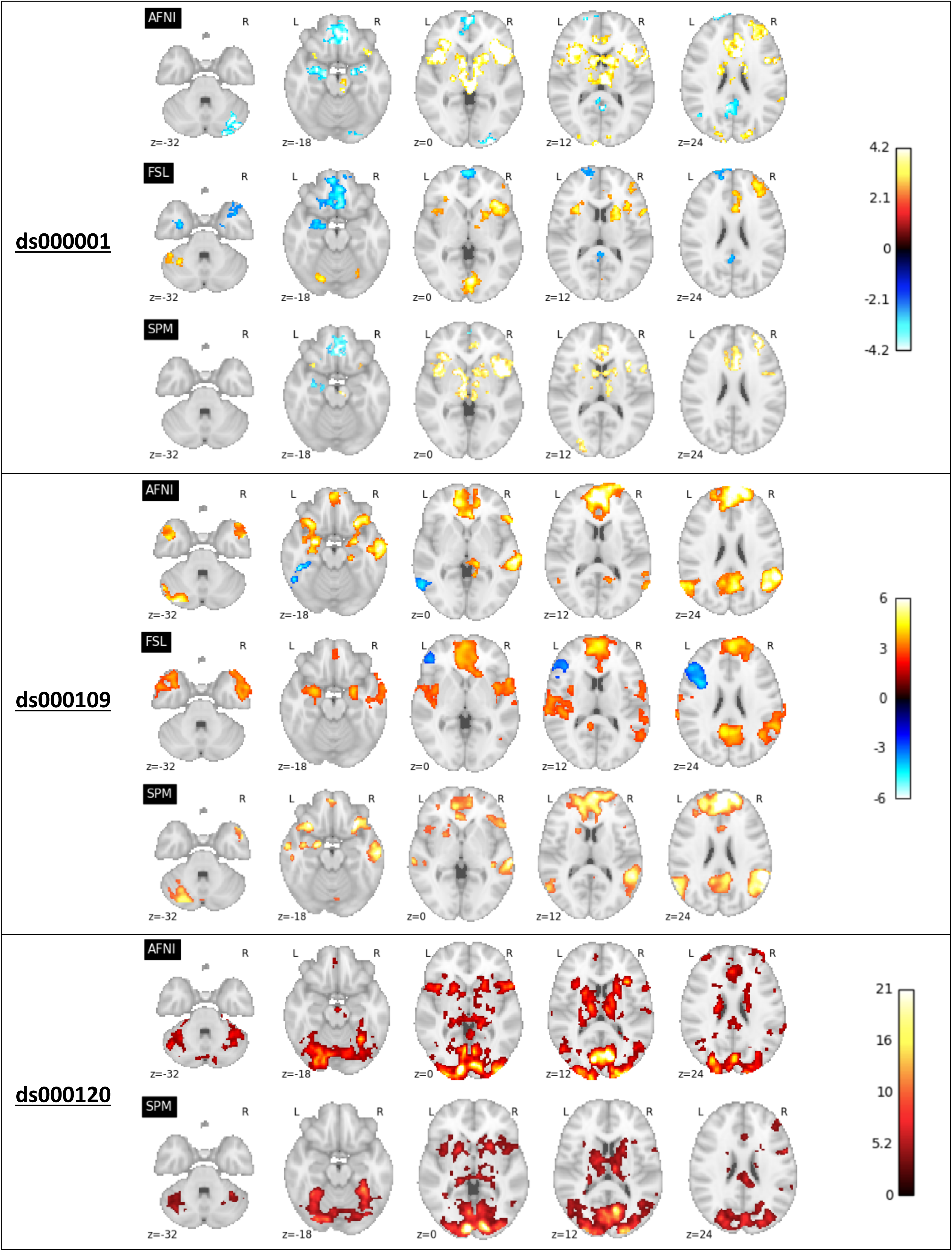
Comparison of the thresholded statistic maps from our reanalysis displayed as a series of axial slices. Top: ds000001’s thresholded T-statistic maps contrasting parametric modulations of the reward balloons versus pumps of the control balloons. Middle: ds000109’s thresholded T-statistic maps of the false belief vs false photo contrast. Bottom: ds000120’s thresholded F-statistic maps of the main effect of time contrast. This figure complements the single slice views shown in Figure 1a.

The ds000109 study reported activations in the bilateral temporoparietal junction. (TPJ), precuneus, anterior superior temporal sulcus (aSTS), and dorsal medial prefrontal cortex (dmPFC). Similar activations from our reanalyses are seen in Figure 1a, middle, although FSL only found activation in the right TPJ and aSTS. Further comparisons shown in Figure 1b, middle, highlight disagreement in the results: AFNI and FSL detected significant deactivations in distinct brain regions (inferior temporal gyrus for AFNI, inferior frontal gyrus for FSL), while SPM did not determine *any* significant deactivation. FSL also found a positive response in the superior temporal gyrus (STG) where AFNI and SPM did not (Fig. 1b, middle, z=0 and z=12 slices).

The original ds000120 study found extensive activations for the main effect of time - the frontal, supplementary, posterior parietal cortex, basal ganglia, prefrontal cortex, ventral striatum and orbitofrontal cortex all showed significant activation. Our reanalyses (Fig. 1a, right) are consistent with these findings, with the exception that neither AFNI nor SPM exhibited orbitofrontal (OFC) activation (though, the SPM analysis mask had poor OFC coverage). AFNI’s F-statistic values look to be generally larger than SPM here (Fig. 1b, bottom, z=0 and z=12 slices). The unthresholded statistic maps from our reanalyses (Figs. 2, S7, S9 & S11) also show that while extreme values display moderate agreement, there are considerable differences across the brain in each given study.

**Figure 2.**
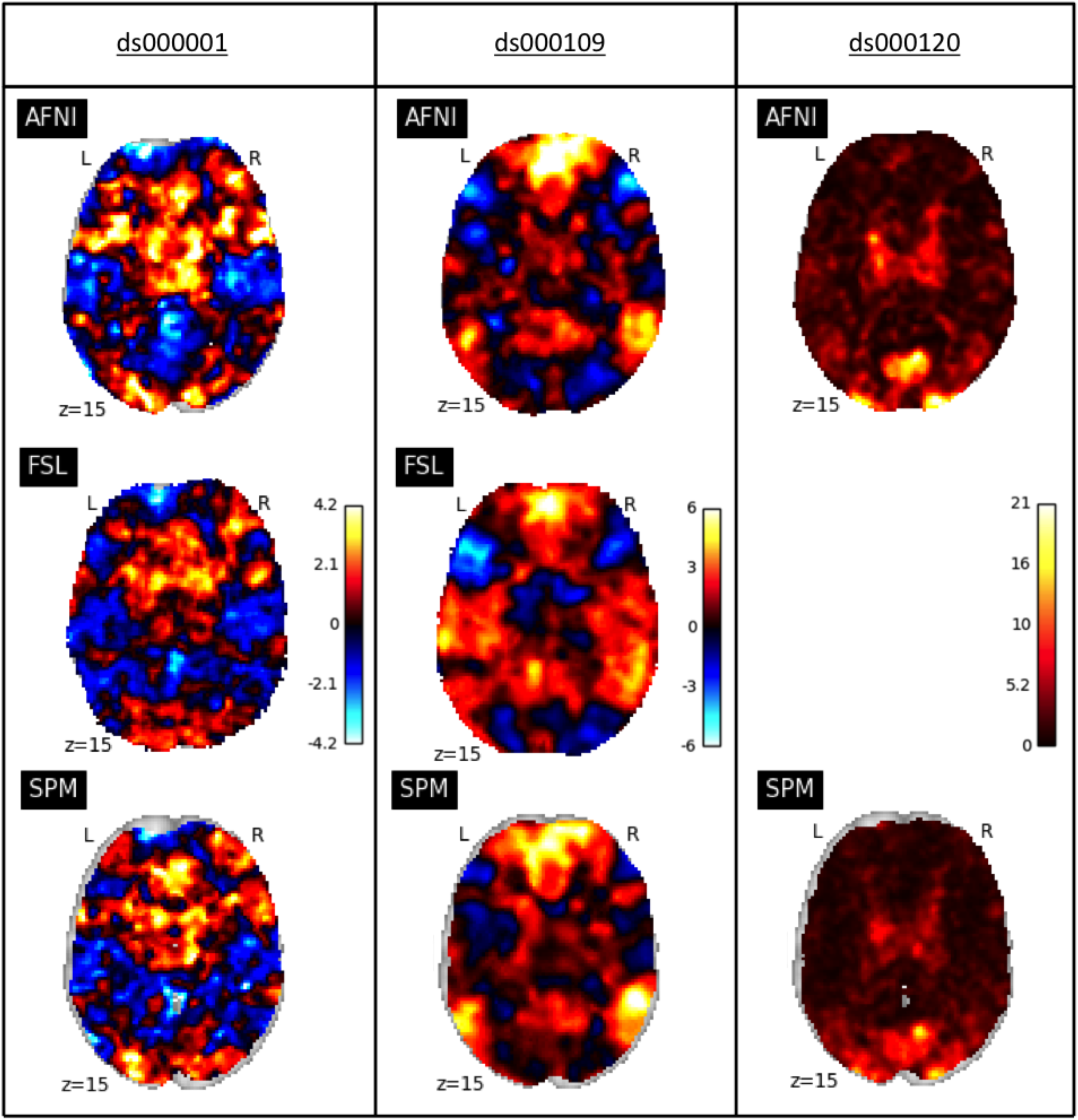
Comparison of the unthresholded statistic maps from our reanalysis of the three studies within each software package. Left: ds000001’s unthresholded T-statistic maps of the parametric modulation of pumps of reward balloons versus the parametric modulation of the control balloon contrast. Middle: ds000109’s unthresholded T-statistic maps of the false belief vs false photo contrast. Right: ds000120’s unthresholded F-statistic maps of the main effect of time contrast. While areas of strong activation are somewhat consistent across all three sets of reanalyses, there is substantial variation in non-extreme values.

NeuroSynth association analyses conducted on the unthresholded T-statistic maps (Table 3) show that the most strongly related term to the activation patterns displayed across all three sets of results was the same: ‘anterior insula’ for each software’s ds000001 map, ‘medial prefrontal’ for ds000109, and ‘visual’ for ds000120. Phrases related to the task paradigm used in each study (‘goal’ for ds000001, ‘theory mind’ for ds000109, ‘visual’ for ds000120) were found across all software’s activation patterns, alongside a range of common anatomical terms.

Figure 3a. compares statistic values across packages using Bland-Altman plots (rendered as 2D histograms) for ds000001 and ds000109. The distribution of the pairwise differences in T-statistics (y-axes) is generally centered about zero, indicating no particular bias, however there is substantial variation here, with T-statistic differences exceeding 4.0 in magnitude. Pairwise correlations ranged from 0.429 to 0.747 for inter-software comparisons (Table 2). The Bland-Altman plots comparing percentage BOLD change maps (Fig. S13) are more conclusive, showing a clear trend for SPM to report larger effect estimates than the other two packages. Figure 3b presents the Bland-Altman plot comparing unthresholded F-statistic images for ds000120, which has a very different appearance due to F-statistics being non-negative. The corresponding Bland-Altman plot comparing partial R^2^ values (Fig. S14) for this study is similar in shape. Broadly speaking, while there are no gross differences in sensitivity, there is a slight tendency for AFNI’s extreme statistics to exceed FSL’s and SPM’s, and SPM’s to exceed FSL’s, most evident in ds000109.

**Table 2.** Summary of Test Statistics. Mean differences and correlations for each pair of test statistic images; mean differences correspond to the y-axes of the Bland Altman plots displayed in Figures 3a, 3b, 7 & 8. Each mean difference is the first item minus second; e.g. AFNI vs. FSL mean difference is AFNI-FSL. Correlation is the Pearson’s r between the test statistic values for the pair compared. Inter-software differences are greater than intra-software.

|  |  | <u>ds000001</u> |  | <u>ds000109</u> |  | <u>ds000120</u> |  |
| --- | --- | --- | --- | --- | --- | --- | --- |
|  |  | **Mean**<br>**Diff** | **Corr** | **Mean**<br>**Diff** | **Corr** | **Mean**<br>**Diff** | **Corr** |
| **AFNI vs. FSL** | Parametric<br>Non-Parametric | 0.009<br>0.271 | 0.616<br>0.577 | 0.035<br>0.006 | 0.585<br>0.573 |  |  |
| **AFNI vs. SPM** | Parametric<br>Non-Parametric | 0.061<br>-0.096 | 0.614<br>0.628 | -0.490<br>-0.445 | 0.747<br>0.787 | 0.415<br>n/a | 0.748<br>n/a |
| **FSL vs. SPM** | Parametric<br>Non-Parametric | -0.047<br>-0.479 | 0.684<br>0.720 | -0.529<br>-0.439 | 0.429<br>0.438 |  |  |
| **AFNI** | Para. vs. NonP. | 0.155 | 0.984 | -0.048 | 0.981 |  |  |
| **FSL** | Para. vs. NonP. | 0.382 | 0.844 | -0.064 | 0.946 |  |  |
| **SPM** | Para. vs. NonP. | 0.000 | 1.000 | 0.000 | 1.000 |  |  |

**Table 3.**
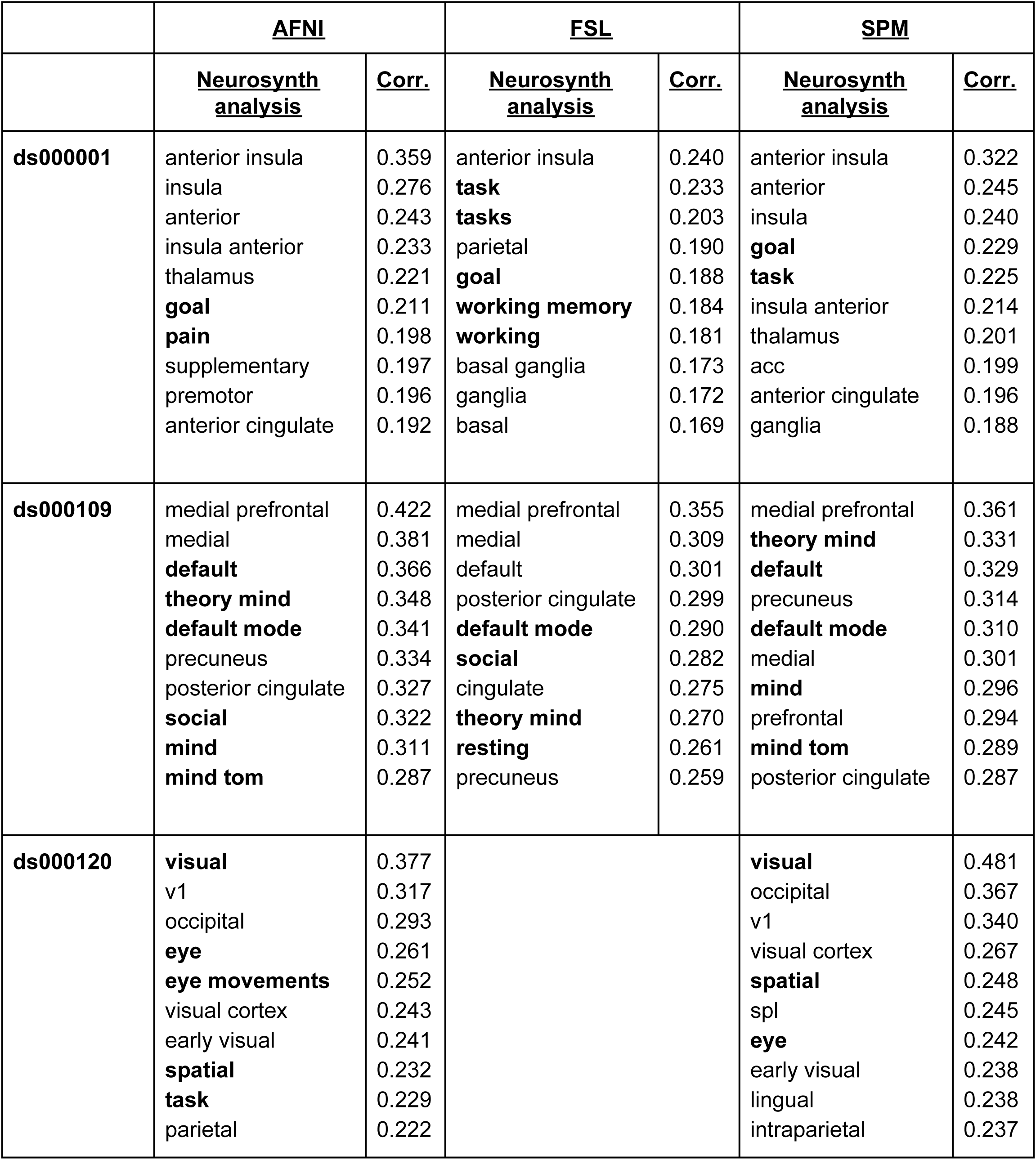
Neurosynth Analyses. The Neurosynth analysis terms most strongly associated (via Pearson Correlation) to each software’s group-level statistic map across the three studies. Non-anatomical terms are shown in bold.

**Figure 3a.**
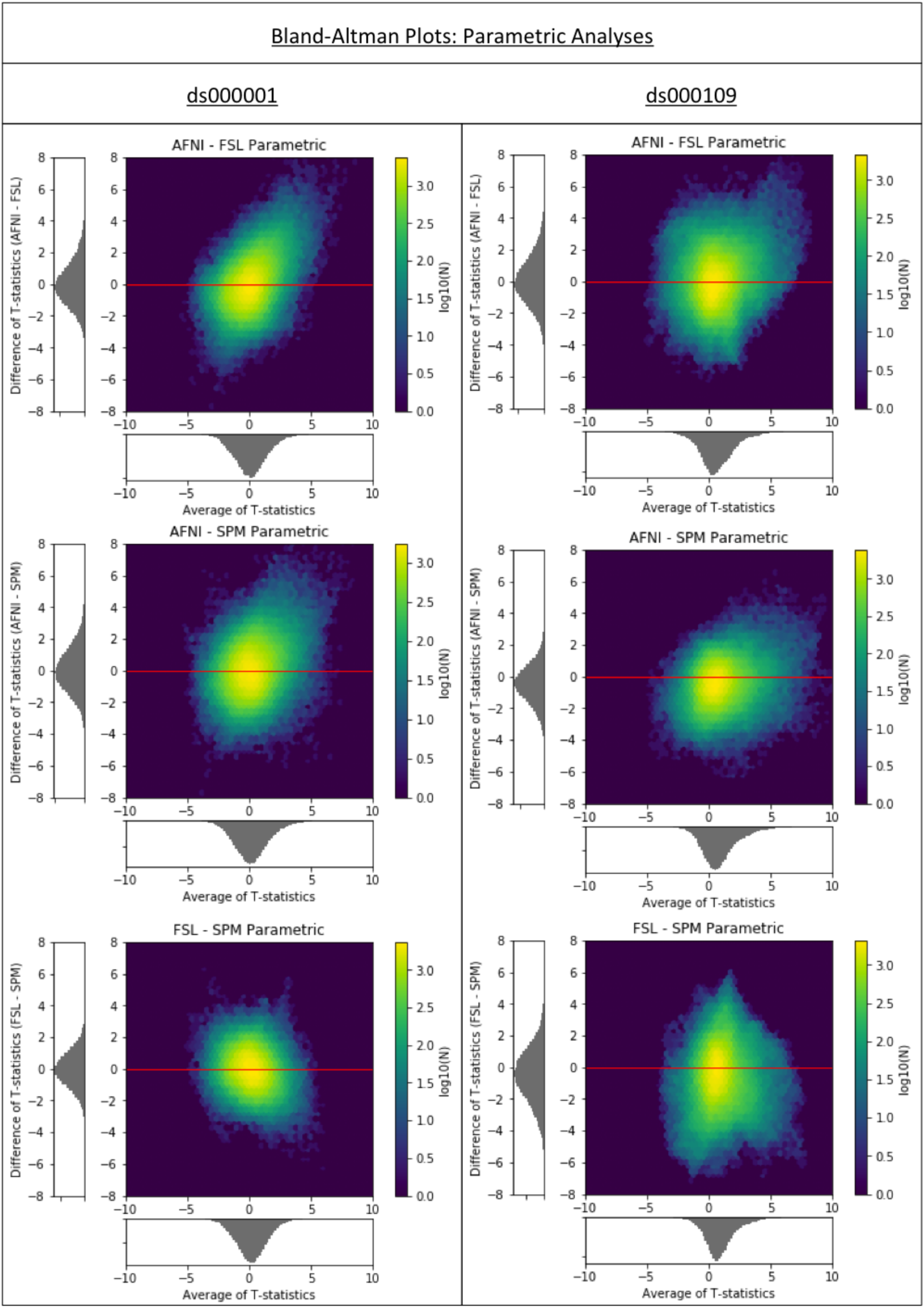
Cross-software Bland-Altman 2D histograms comparing the unthresholded group-level T-statistic maps computed as part our reanalyses of the ds000001 and ds000109 studies within AFNI, FSL and SPM. Left; Comparisons for ds000001’s balloon analog risk task, T-statistic images contrasting the parametric modulation of pumps of the reward balloons versus parametric modulation of pumps of the control balloon. Right; Comparisons for ds000109’s false belief task, T-statistic images contrasting the false belief versus false photo conditions. Density images show the relationship between the average T-statistic value (abscissa) and difference of T-statistic values (ordinate) at corresponding voxels in the unthresholded T-statistic images for each pairwise combination of software packages. While there is no particular pattern of bias, as the T-statistic differences are centered about zero, there is remarkable range, with differences exceeding +/-4 in all comparisons.

**Figure 3b.**
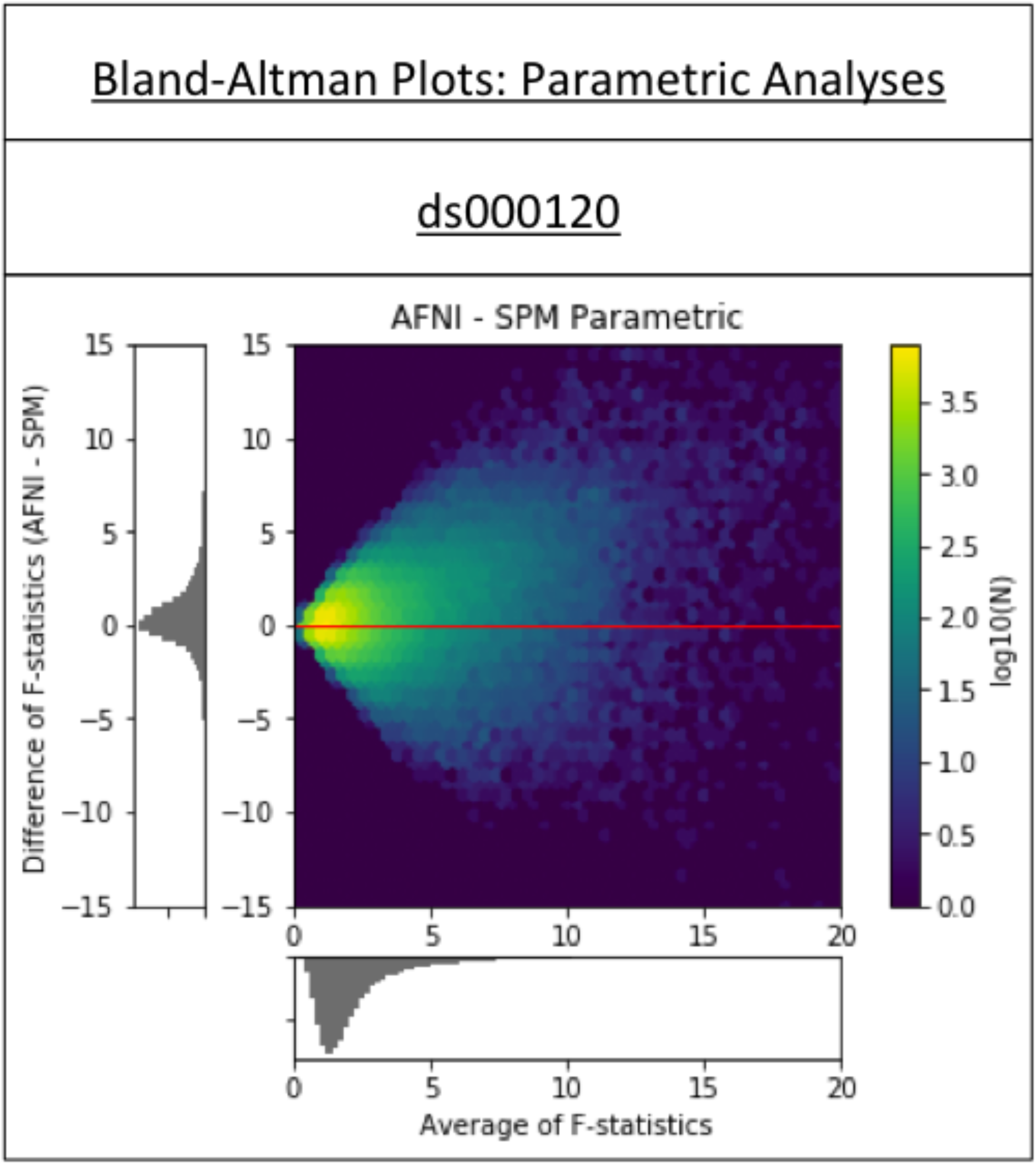
Cross-software Bland-Altman 2D histogram comparing the unthresholded main effect of time F-statistic maps computed in AFNI and SPM for reanalyses of the ds000120 study. The differences are generally centered about zero, with a trend of large F-statistics for AFNI. (The funnel-like pattern is a consequence of the F-statistic taking on only positive values.)

Spatial localization of significant activation in the thresholded T-statistic images also varied across software packages. Figure 4 shows the Dice coefficients for all pairs of analyses (parametric results are presented in first 3 rows of larger triangles). For ds000001, the average value of Dice coefficients comparing locations of activations across reanalyses is 0.379. These values improve for ds000109, where the mean Dice coefficient for positive activations is 0.512. Here, AFNI and FSL were the only software packages to report significant negative clusters for the ds000109 study. Strikingly, these activations were found in completely different anatomical regions for each package, witnessed by the negative activation AFNI/FSL dice coefficient of 0. Finally, the AFNI/SPM Dice coefficient for the thresholded F-statistic images obtained for ds000120 is 0.684; it is notable that across all studies, the AFNI/SPM dice coefficients are consistently the largest.

**Figure 4.**
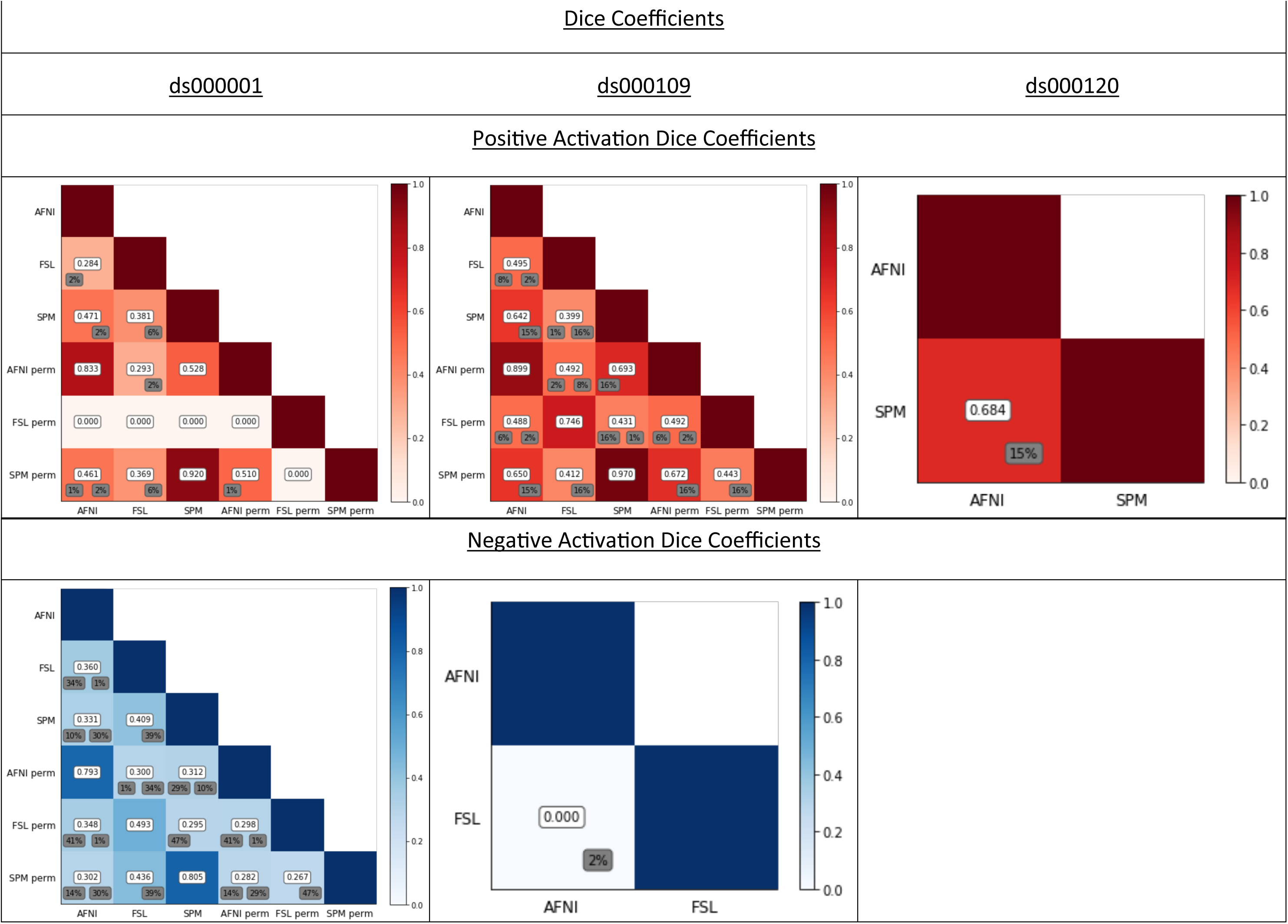
Dice coefficients comparing the thresholded positive and negative T-statistic maps computed for each pair of software package and inference method for each of the three reproduced studies. Dice coefficients were computed over the intersection of the pair of analysis masks, to assess only regions where activation could occur in both packages. Percentage of ‘spill over’ activation, i.e. the percentage of activation in one software’s thresholded statistic map that fell outside of the analysis mask of the other software is displayed in grey; left value for row software, right value for column software. For ds000001 increases, FSL permutation obtained no significant results, thus generating Dice coefficients of zero; for ds000109 decreases, only AFNI and FSL parametric obtained a result and hence only one coefficient is displayed. Dice coefficients are mostly below 0.5, parametric-nonparametric intra-software results are generally higher; ds000120’s F-statistic results are notably high, at 0.684, perhaps because it is testing a main effect with ample power.

Spill over values, given by the grey values beneath the dice coefficients in Figure 4, are generally largest for SPM comparisons. They are particularly prominent in the negative activation plot for ds000001, where there is at least 30% spill over for all parametric pairwise comparisons, the largest being 39% for SPM/FSL. Recalling that these values are the percentage of activation which occured within one package that was outside the other package’s analysis mask, this is likely due to the fact SPM consistently had the smallest analysis mask out of the three packages, while FSL had the largest. In our ds000001 reanalyses, SPM’s group-level analysis mask was made up of 175269 voxels, while AFNI’s had 198295 voxels and 251517 for FSL. For ds000109, SPM’s group-level mask contained 178461 voxels compared to AFNI’s 212721 and FSL’s 236889. Finally, for ds000120, SPM had 174059 voxels to AFNI’s 208340. (Note FSL’s mask image has slightly but consistently more non-zero voxels than in its statistical result images).

Further evidence of high spatial variability is also exhibited by the Euler Characteristic plots for the parametric analyses presented in Figure 5a, top (and supplementary Fig. S12 for ds000120), complemented by the cluster count plots in Figure 5b, top. Since the Euler Characteristic counts the number of clusters minus the number of ‘handles’ plus the numbers of ‘holes’ in an image, for large thresholds we expect the Euler Characteristic to closely approximate the number of clusters of significant activation present in the equivalent thresholded map. This is confirmed by Figures 5a and 5b, both figures showing that across the two studies FSL had a smaller number of activated clusters at larger thresholds. For ds000001, the peak cluster count value (Figure 5b, top left) occurs at a lower threshold for FSL. This plot suggests that in general FSL’s T-statistic values were more liberal here - the initial rise of the FSL curve signifies the T-statistic image breaking up into clusters at lower thresholds than AFNI and SPM, and then as the clusters begin to get ‘thresholded out’ this causes the FSL curve to dip below the other two packages. The Euler Characteristic plots instead highlight overall topological differences in the statistic maps: If the images were the same up to an image-wide monotonic transformation, this would be revealed by the Euler Characteristic curves having the same general shape but with some portions shifted or compressed. While the ds000001 curves are more similar in shape, ds000109’s curves are notable for each having a highly distinct shape, indicating substantially different topologies of activation.

**Figure 5a.**
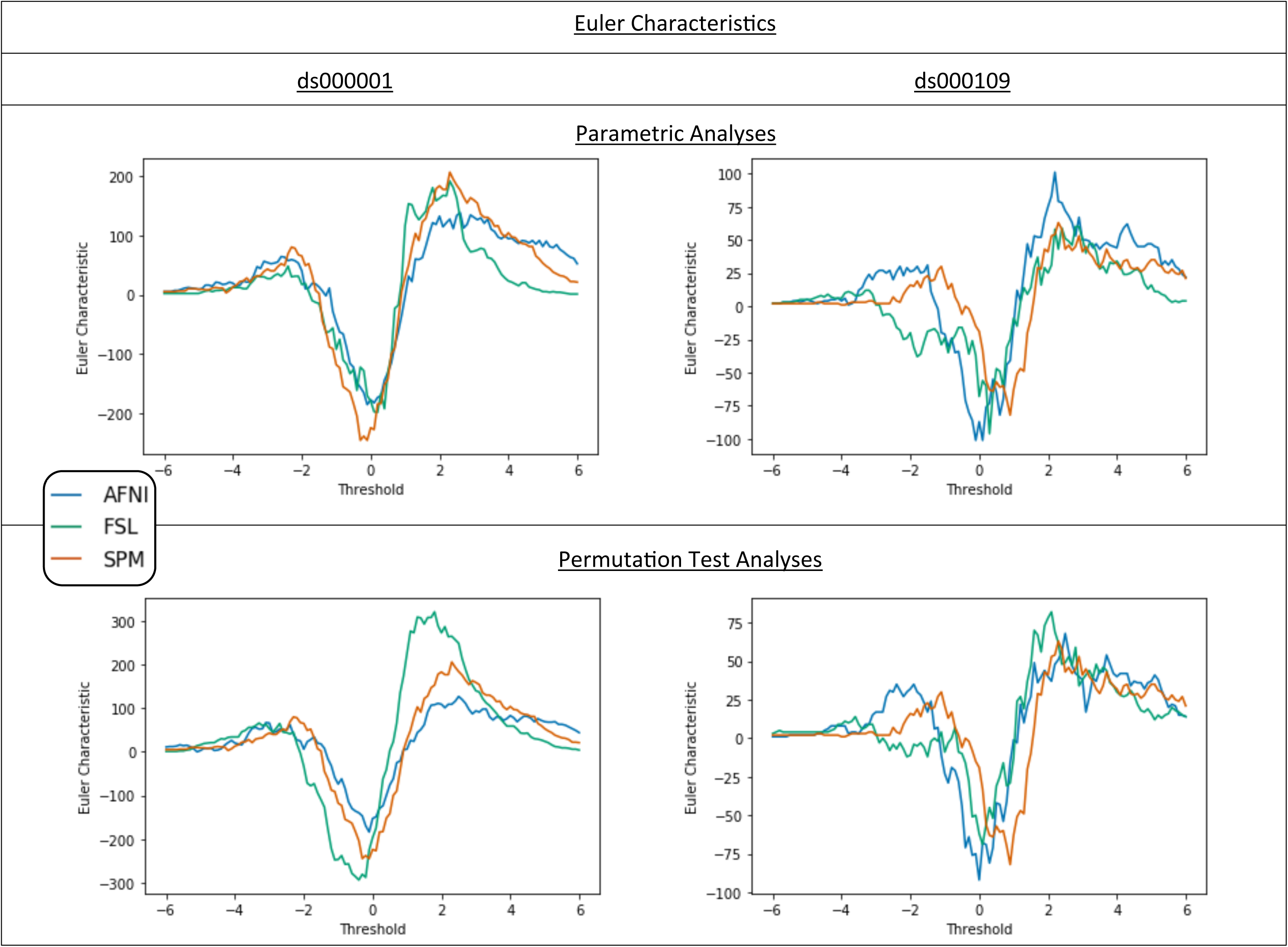
Euler Characteristic plots for ds000001 and ds000109. On top, comparisons of the Euler Characteristic computed for each software’s T-statistic map from our reanalyses using a range of T-value thresholds between -6 and 6. Below, comparisons of the Euler Characteristics calculated using the same thresholds on the corresponding T-statistic images for permutation inference within each package. For each T-value the Euler Characteristic summarizes the topology of the thresholded image, and the curves provide a signature of the structure of the entire image. For extreme thresholds the Euler Characteristic approximates the number of clusters, allowing a simple interpretation of the curves: for example, for ds000001 parametric analyses, FSL clearly has the fewest clusters for positive thresholds.

**Figure 5b.**
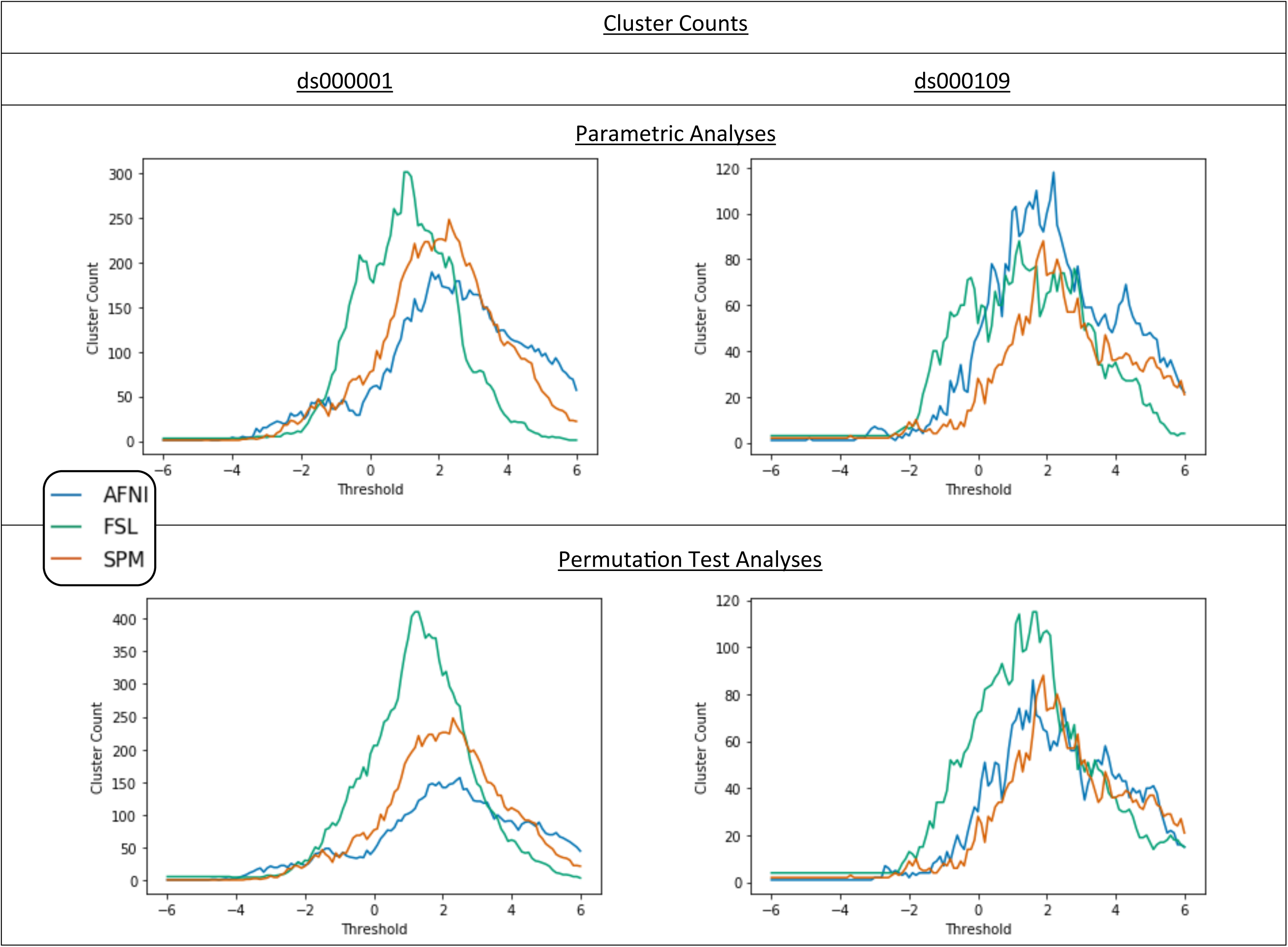
Cluster count plots for ds000001 and ds000109. On top, comparisons of the number of cluster found in each software’s T-statistic map from our reanalyses using a range of T-value thresholds between -6 and 6. Below, comparisons of the cluster counts calculated using the same thresholds on the corresponding T-statistic images for permutation inference within each package.

### 3.2 Cross-Software Variability for Nonparamteric Inference

Consistent with the parametric inference results, activation localization and statistic values varied greatly between packages for the permutation test results computed for ds000001 and ds000109.

Before reviewing statistic map comparisons, we stress that the goal of these nonparametric analyses is to obtain FWE-corrected cluster p-values with weaker assumptions. Thus the permutation test unthresholded statistic maps are not “nonparametric” maps, but rather usual one-sample T-test maps that form the basis of permutation analyses. While SPM’s parametric analysis uses the same one-sample T-test, AFNI’s and FSL’s parametric models use a mixed-effects model and weighted least squares. Hence all comparisons of the nonparametric test statistic values (in contrast to thresholded maps) do not convey information about nonparametric inference per se, but compare different preprocessing and first level modeling from the three packages while holding the second level model constant.

Quantitative assessment with Dice coefficients are shown in Figure 4 (“perm” vs “perm” cells) and - in accordance with the parametric results - are generally poor. Like the parametric analyses, AFNI/SPM Dice values are altogether better than the other comparisons. For ds000001, FSL’s nonparametric method found no significant clusters, and thus all Dice coefficients connected to this analysis are zero. However, note that the significant regions found in the other parametric and nonparametric results for this study mostly comprise of a single activation cluster spanning the lateral and medial frontal cortex, insular cortex, basal ganglia, and brainstem - an extensive and irregularly-shaped cluster that could easily become disconnected and thus lose significance. As before, ds000109 Dice values are also generally better than ds000001.

The nonparametric Bland-Altman plots (Fig. 6) show substantial spread qualitatively similar to the parametric ones (Fig. 3a), and correlations between statistics maps are similar for nonparametric in congruence with the parametric comparisons (Table 2). EC curves (Fig. 5, bottom) again exhibit considerable topological variation between software packages. Notably, while AFNI and SPMs EC curves are relatively similar across choice of inference method, FSL permutation inference determined substantially more clusters than parametric for low positive thresholds in both studies (Figs. 5a and 5b, bottom).

**Figure 6.**
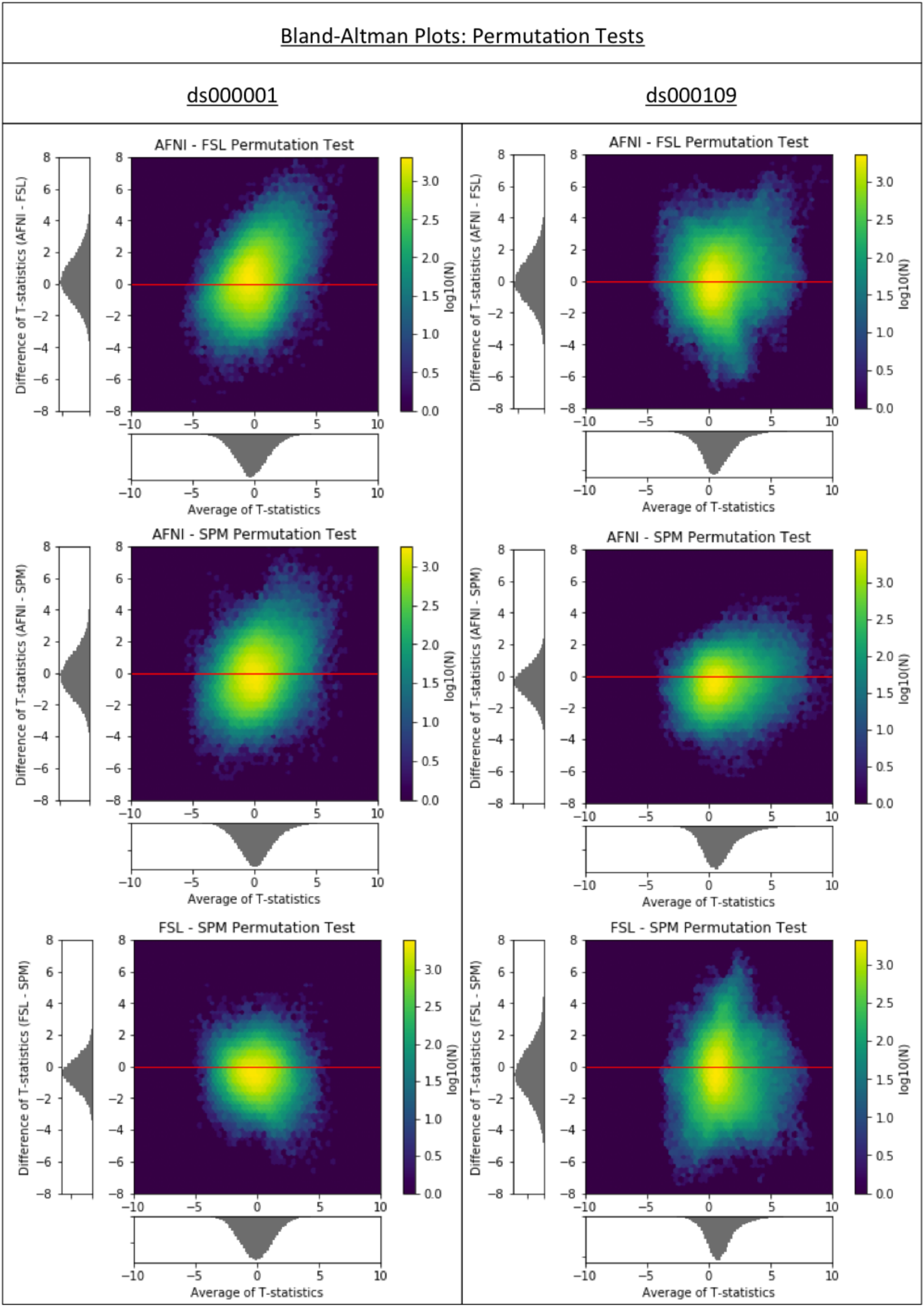
Cross-software Bland-Altman 2D histograms for the ds000001 and ds000109 studies comparing the unthresholded group-level T-statistic maps computed using permutation inference methods within AFNI, FSL and SPM. Similar to the results obtained using parametric inferences in Figure 3a, all of the densities indicate large differences in the size of activations determined within each package.

### 3.3 Intra-Software Variability, Parametric vs nonparamteric

Comparisons of parametric and permutation test inference results within each package hold all preprocessing and first level modelling constant, only varying the second level model and inference procedure. The level of agreement between the two inferences *within* each package varied greatly across software. Before making comparisons, we note that since SPM’s parametric and nonparametric inference share the same group level model, the unthresholded statistic images produced using each inference model are identical here.

The thresholded statistic maps are generally similar within each of the software packages (ds000001: Fig. S2 vs Fig. S3; ds000109: Fig. S4 vs S5), with the exception of FSL’s nonparametric inference ‘decreases-only’ finding for ds000001. Unthresholded maps are notably more similar for ds000109 (Fig. S9 vs S10) than for ds000001 (Fig. S7 vs Fig. S8), again noting that SPM’s pairs of maps here are identical.

Bland-Altman plots (Fig. 7) reveal much greater levels of parametric-nonparametric agreement, with AFNI displaying greater agreement than FSL. For FSL, we selectively investigated voxels that differed by the greatest amount, and often found individual subjects responsible: A single subject with a large observation can drive a conventional one-sample T-test, but when that same subject also has large intrasubject variance FSL’s mixed effect model downweights that subject leading to a substantially different T-test. The increased difference in AFNI’s values for ds000109 for larger statistic values could also reflect a similar downweighting procedure within the software.

**Figure 7.**
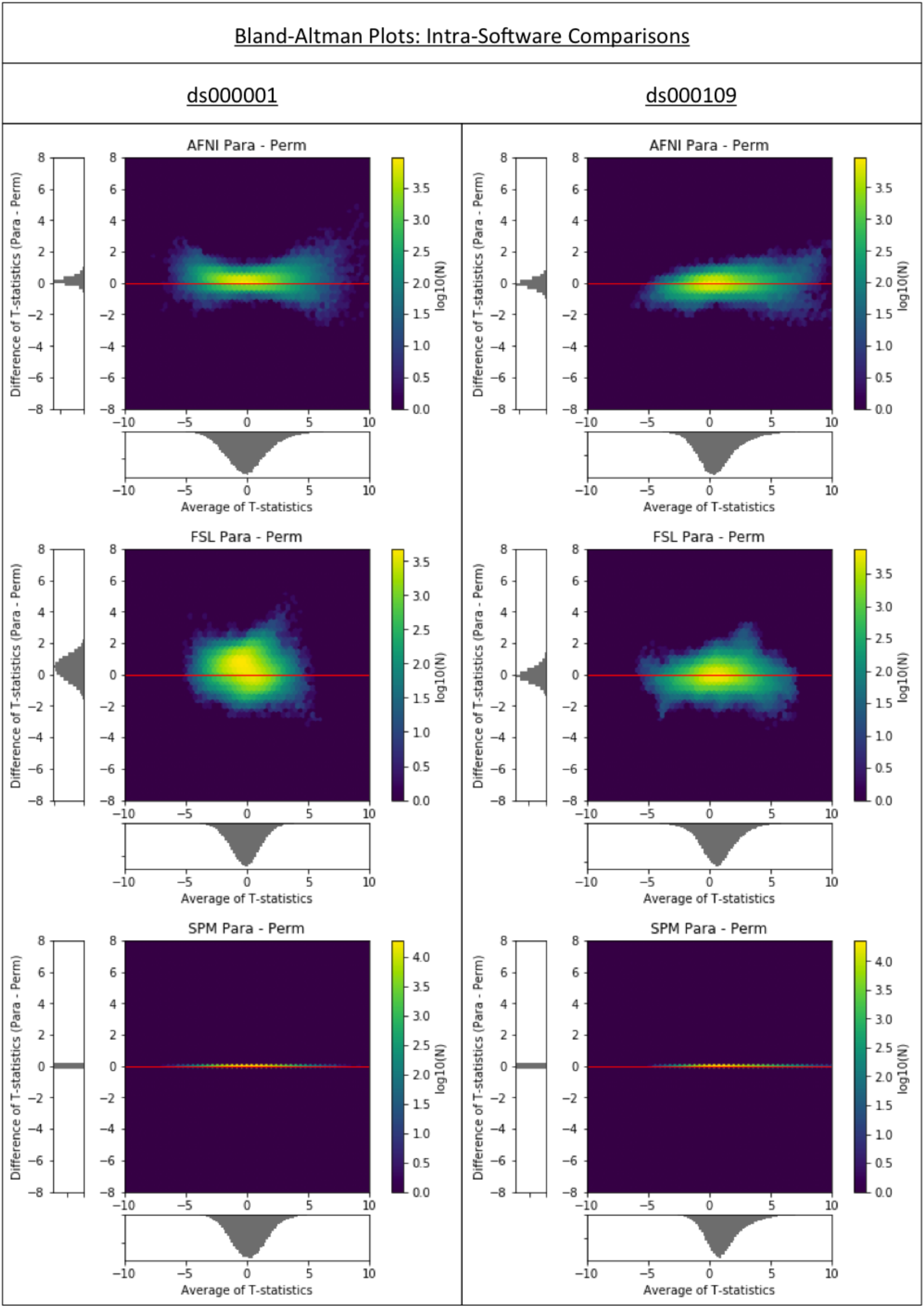
Intra-software Bland-Altman 2D histograms for the ds000001 and ds000109 studies comparing the unthresholded group-level T-statistic maps computed for parametric and nonparametric inference methods in AFNI, FSL and SPM. Each comparison here uses the same preprocessed data, varying only the second level statistical model. SPM’s parametric and nonparametric both use the same (unweighted) one-sample T-test, and thus show no differences. AFNI and FSL’s parametric models use iterative estimation of between subject variance and weighted least squares and thus show some differences, but still smaller than between-software comparisons.

The Dice coefficients comparing the thresholded permutation test and parametric inferences are generally the best of any (Fig. 4, 3-element lower diagonal). In general the origin of parametric-nonparametric differences are parametric inference finding a slightly larger number of clusters significant.

### 3.4 Conclusion

Across all three of the studies reanalysed here we have discovered considerable differences between the AFNI, FSL, and SPM results. The scale of these differences has been highlighted by each of the quantitative metrics applied to compare the group-level statistic maps: Dice coefficients were commonly less than 50% for cross-software comparisons, Bland-Altman plots showed that differences between reported T-statistic values were as large as 4 for a considerable quantity of voxels, and Euler Characteristic curves displayed a divergence in the number of clusters being reported in each software - even at large thresholds.

In reporting these comparisons, we are not making any statements as to which software package is better or worse. Without a gold standard to compare to, we do not believe that such an endeavour would be fruitful. Rather, we feel that the key contribution of our work is the quantitative measurement of inter-software differences on common datasets. Our finding that exceedingly weak effects may not generalise across packages - evidenced across all three of our analyses - is the primary take-home message of this work. While larger effects were found to be more robust, we stress that our analyses have been conducted under particularly favourable conditions: the use of studies with a strong, primary effect and extensive efforts made to harmonise the three analyses. Because of this, at best our results present an optimistic view of inter-software disparities. To better understand the underlying differences between software, further work on quantification of pipeline-related variation is needed, which will hopefully lead to harmonisation in software implementation to reduce these differences in the future. Another line of work would be the creation of integrative intra-study, ensemble learning techniques to integrate inconsistent findings. An additional contribution with this effort is to provide generalizable measures and metrics to enable software validation, which we hope may benefit any further comprehensive comparison of software packages.

## 4. Discussion

Our results have displayed extensive differences in the results between the analysis pipelines of the three software packages. The low Dice values and differences in Euler characteristics (Figs. 4 & 5a) are particularly salient, showing heterogeneity in the sizes and shapes of clusters determined across packages, indicating a strong dependance on software in terms of the anatomical regions covered by the activation. While some authors have pointed out the limitations of interpreting differences in the set of voxels that make up a significant activation when using clusterwise inference (which makes assessments based on topological properties of statistical maps) (Chumbley and Friston, 2009), we see merit in the use of quantitative measurements such as Dice due to the ultimate application of statistical maps to infer on precise areas of the brain active during a task. While a deeper analysis on the differences between topological aspects of images across software would be valuable, there are inherent difficulties in this approach, such as identifying corresponding features between maps when the number and size of activations reported are variable across software.

It is notable that the level of variation in our analysis results also fluctuated across the datasets we analyzed. This is highlighted in our Dice comparisons, where the ds000001 Dice coefficients are considerably smaller than ds000109 for both the inter and intra-software comparisons. The relatively poor performance of ds000001 may be due to the smaller sample size for this study (16 vs 21 for d000109), as well as the particular inference method used. For ds000001, group-level inference was conducted using a cluster-forming threshold of p < 0.01 uncorrected. A recent study (*Eklund et al., 2016*) found that parametric inference for a one-sample T-test at this threshold in AFNI, FSL and SPM resulted in false-positive rates far exceeding the nominal level - severely for cluster-forming threshold p < 0.01, modestly for p < 0.001 - while nonparametric permutation performed closer to the expected 5% FWE level. The results obtained here for ds000001 are consistent with these findings: across all three software packages, the thresholded images produced from permutation test inference display fewer significant clusters than the corresponding parametric maps. While the cluster-defining threshold p < 0.005 applied in the ds000109 study was not analyzed in *Eklund et al.*, consistency between packages using parametric and nonparametric inference was greater for this study.

Notably, while all packages are purportedly using the same MNI atlas space, an appreciable amount of activation detected by AFNI and FSL fell outside of SPM’s analysis mask (shown by the ‘spill over’ values displayed in gray, Fig. 4). Considerable differences in mask sizes are likely to have been a major factor for the disparities in activation and low dice coefficients seen across packages. For effects close to the edge of the brain, a larger analysis mask allows for a larger cluster volume, which can ultimately be the difference as to whether a cluster is determined as significantly active or not. This may explain why only FSL - which had the largest analysis mask - found an auditory response in the ds000109 study, or why Dice coefficients are generally worse for negative activations than positive in our ds000001 renanalyses, where positive clusters were on-the-whole reported in more central anatomical regions. Another possible reason for poor Dice values here is that the size and number of clusters determined for negative activations was smaller than that of positive activations. Since Dice and spill-over values are proportional measurements, this means they will have been more susceptible to differences in cluster and mask size for the negative activations relative to the positive. Disagreement in atlas space may have contributed to the lack of structure in the Bland-Altman plots, however no gross misalignment between packages was evident (Fig. S1). While far from perfect, the ds000120 AFNI and SPM thresholded results have the best Dice similarity score, likely due to the use of a very strong main effect as an outcome of interest.

Qualitative comparison of the results provide some optimism, with certain patterns of activation found across all packages. For example, the ds000001 parametric analyses were unanimous in determining significant activation in the anterior insula. While there is greater discordance over the precise location of activation within the anterior insular region, as well as the precise statistic values here, altogether our results align. This may substantiate that the strongest effects are robust across packages, supported by our own comparisons of the unthresholded maps that showed moderate agreement between software packages in areas with strong signal (many of these anatomical regions were identified across all three package’s NeuroSynth association analyses) but greater disagreement elsewhere, ds000109 and ds000120 displaying more consistency than ds0000001. However, in making these qualitative comparisons, what has become most transparent is the importance for researchers to - at the very least - share their final statistical maps. The reasons for this are exhibited most clearly by our ds000109 analyses; the visual slice comparison of our replications of the main figure from the original study in Figure 1a, shown alongside the publication figure itself, look remarkably similar and could lead to the conclusion that each package’s results highly agree. It is only when analyzing these results over the whole brain, that we discover broad differences in these activation patterns, e.g. positive activation identified in the auditory cortex in FSL that was neither picked up by AFNI and SPM, and significant deactivation determined only by AFNI and FSL.

Subject data were missing from both the ds000109 and ds000120 datasets. For ds000109, while 29 young adults were scanned for the false belief task, only 21 were present in the dataset; for ds000120, we analyzed 17 subjects instead of 30 used in the original study. These analyses therefore should not be compared like-for-like with the published results, and have substantially less statistical power than the original studies. Overall, our sample sizes for the three datasets analyzed (ds000001, ds000109, ds000120) are 16, 21, and 17 respectively. While small, these sample sizes are fairly representative of a typical functional neuroimaging study over the past two decades - between 1995 and 2015, the median sample size of an fMRI study increased steadily from 8 to 22 (Poldrack et al., 2017). This increase has continued, and a review of 2017 publications found a median sample size of 33 (Yeung 2018). Hence while our datasets are important for judging previous studies, a future comparison exercise with larger datasets would be a valuable addition to the literature.

At the start of our investigations, we selected a common set of preprocessing steps to be applied within each software package across all studies regardless of whether they had been used in the original analysis. This was to maximise the comparability of the results while being consistent with best practices within the community. However, several complications arose during our analyses. For ds000001, orientation information was missing from seven of the subject’s structural and functional scans. Because the source DICOM files were no longer available, it was not possible to retrieve the original position matrices. As a consequence of this, the structural and functional images were misaligned, resulting in suboptimal coregistration during our analyses. Additionally, a bug in the event-files induced during data conversion to the BIDS standard had resulted in some of the event timings being lost. Thanks to the cooperation of BIDS and OpenfMRI these problems were solved; a revised dataset (Revision: 2.0.4) was uploaded to OpenfMRI and used in our analysis.

This study has mainly focused on comparing statistic maps, since these are the images studied to make judgments about localisation and determine the neuroscientific interpretation of results. However, by comparing the statistic maps obtained at the end of the pipeline, we have only assessed the net accumulation of differences introduced at each stage of the analysis procedure. Further work could include a detailed investigation of which steps in the analysis contribute most to this variation. A future study could explore this by considering the factorial expansion of all possible combinations of preprocessing, first level modelling, and second level modelling, akin to previous efforts in assessing reproducibility over a number of pipelines (Strother et al., 2002).

Finally, we acknowledge the central influence that data sharing had on this work. This study would not have been possible without OpenfMRI and the pool of researchers who have contributed their data to the project, to whom we are grateful. Due to the restrictive requirements of this investigation - the necessity for published task-based fMRI data using analysis methods compatible in AFNI, FSL, and SPM - the three studies analyzed here were found to be the only datasets hosted on OpenfMRI suited to the aims of our investigation. Of the datasets that were not used, the most common reasons for exclusion were that no publication was associated to the data, that the sample size of the study was too small, or that custom software or region of interest analysis had been used as part of the analysis pipeline which was not feasible across the three software packages. Nevertheless, a greater sample of studies will need to be replicated across the packages to gain a more comprehensive understanding of the variability between software, with the inclusion of analyses that use small volume corrections that are becoming increasingly common within the field. We have kept many parameters fixed in our analyses, such as the use of non-linear registration for all software packages, and the addition of motion regressors in all our design matrices. How changes in these variables influence the analysis results warrants further investigation; for example, while we decided to fix a 2mm cubic voxel size in all packages (since this is the default in FSL, SPM), a recent study found that alterations in this parameter can significantly impact statistical inference (Mueller et al., 2017). The use of a wider range of software packages (e.g. FreeSurfer; RRID:SCR_001847; (Dale et al., 1999)), as well as different software versions which were not accounted for in the present study would also strengthen any future analysis.

Future efforts would be strengthened by additional sharing of analysis scripts and statistic maps, enabling confirmation of analyses that follow original procedures and permitting more quantitative comparison of statistic maps. We have made all of our analysis scripts available and statistic maps available, and we hope more researchers join this trend to advance openness in neuroimaging science.

## 5. Acknowledgments

This work was supported by the Wellcome Trust, grant 100309/Z/12/Z. We are extremely grateful to the AFNI team for their manuscript (Taylor et al., 2018) that gave extensive feedback on the AFNI analyses we conducted in an earlier preprint version of this work. We would particularly like to thank Rick Reynolds and Daniel Taylor for the help they gave us, as well as further suggestions from Daniel Glen and Dylan Nielson. We thank Krzysztof Gorgolewski for assistance with OpenfMRI/OpenNeuro; and for comments from Stephen Smith and Jesper Andersson on FSL.

