## Supplementary Materials for "Exploring the Impact of Analysis Software on Task fMRI Results" (from another version of this preprint: version 1, 2018-03-20)

### Supplementary Figures

Figure S1. Registration QC: Mean and standard deviation of anatomical and mean functional images

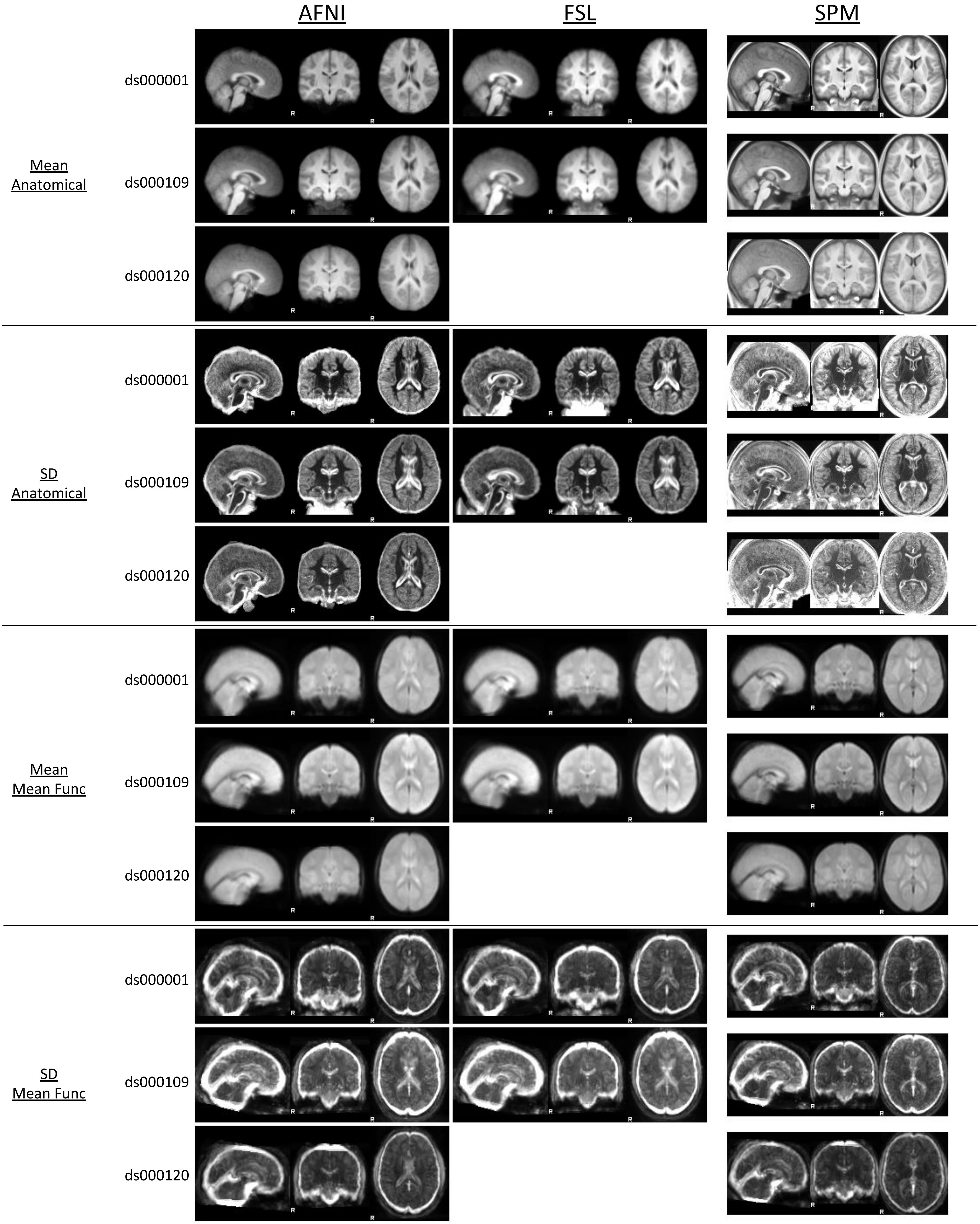

Figure S2. ds000001 Inter-Software Comparison, 5% FWE Clusterwise Inference

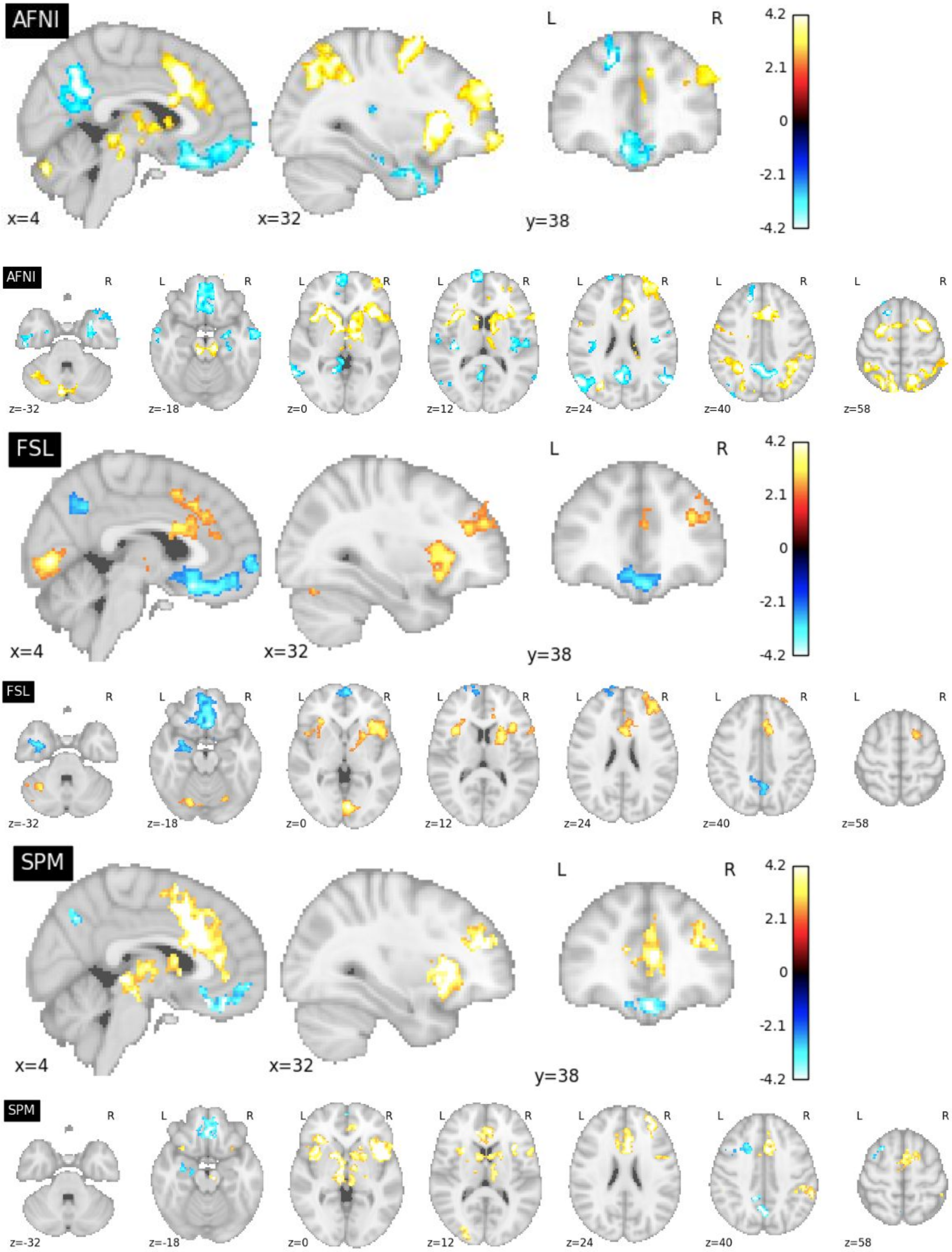

Figure S3. ds000001 Inter-Software Comparison, 5% FWE Clusterwise Permutation Inference

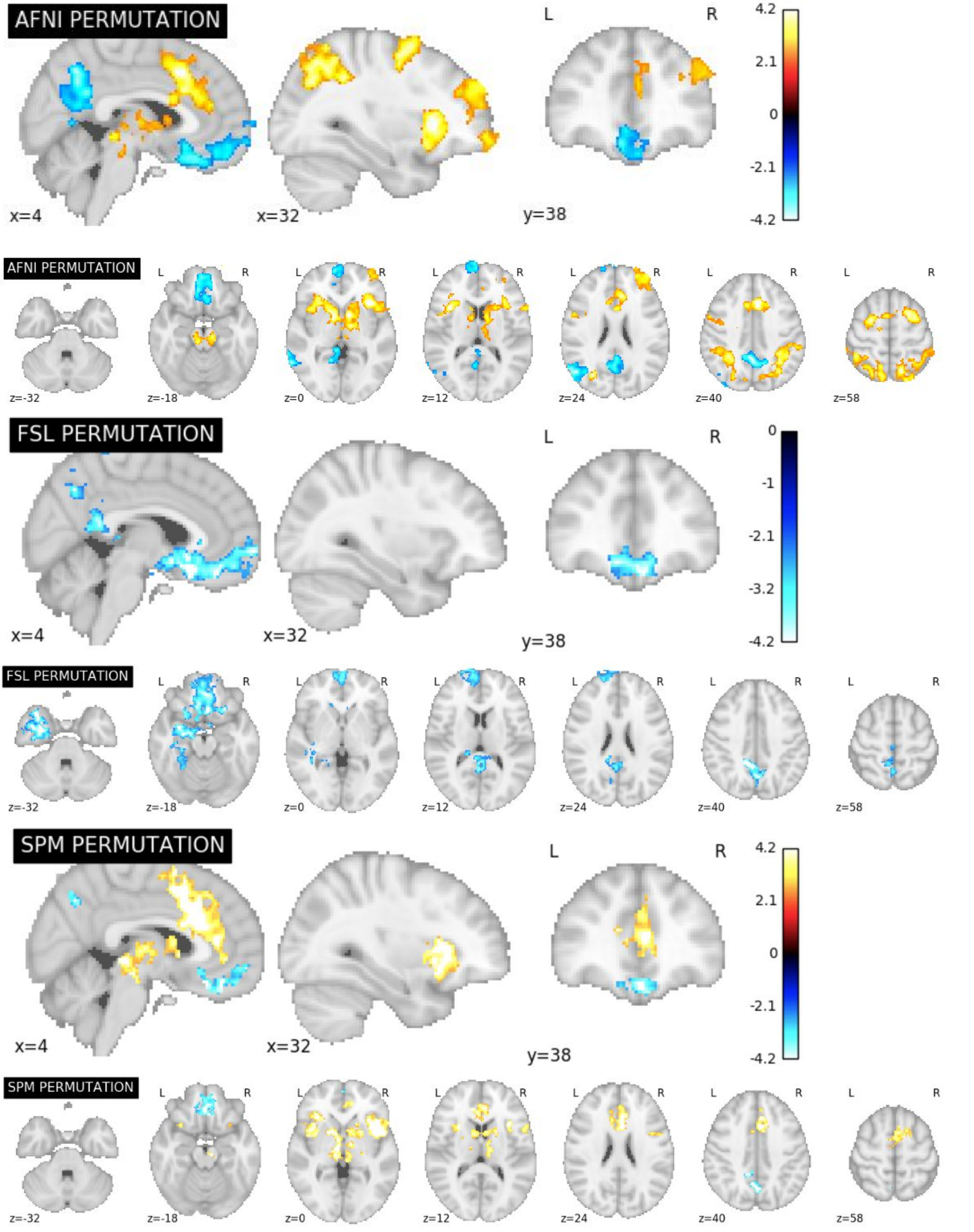

Figure S4. ds000109 Inter-Software Comparison, 5% FWE Clusterwise Inference

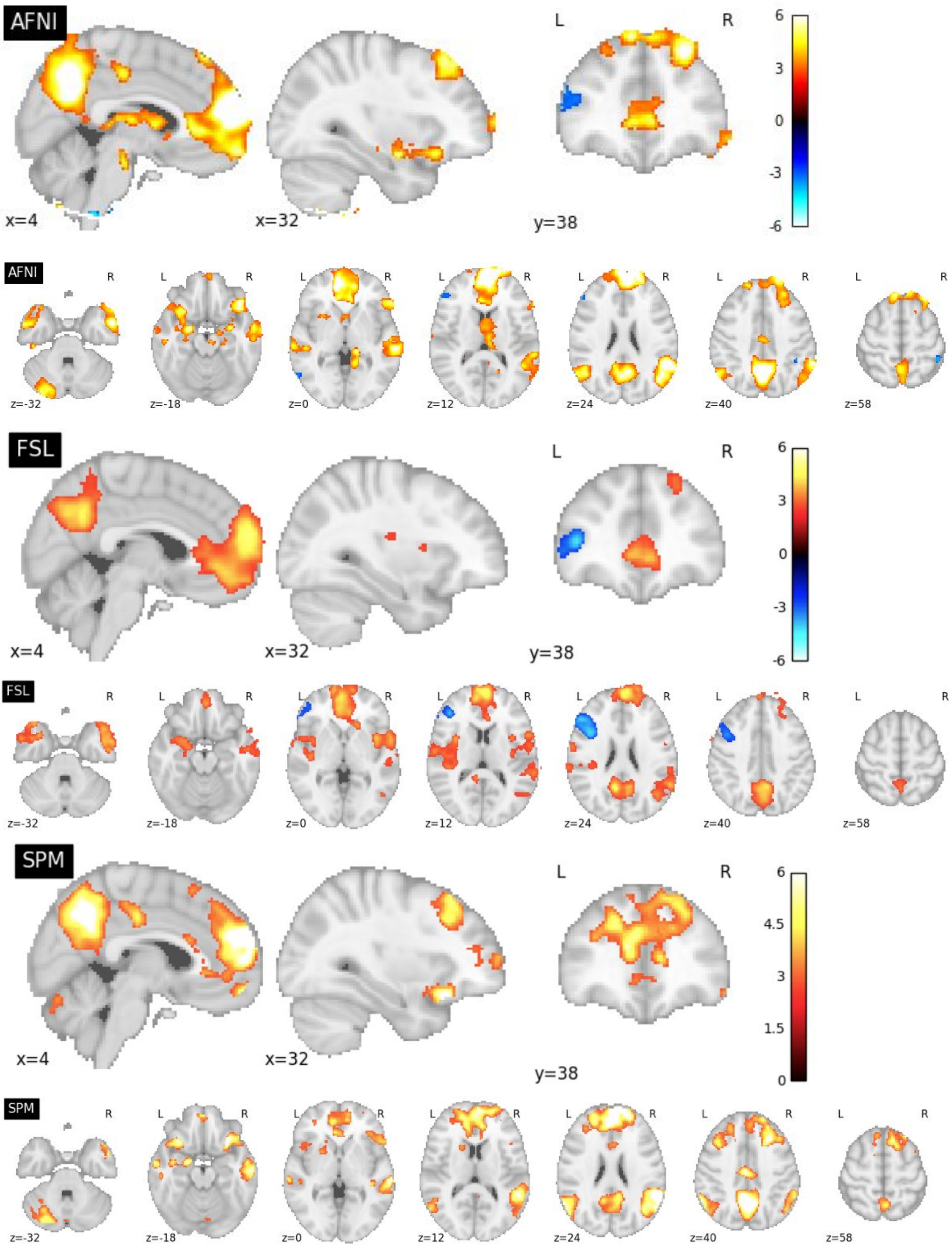

Figure S5. ds000109 Inter-Software Comparison, 5% FWE Clusterwise Permutation Inference

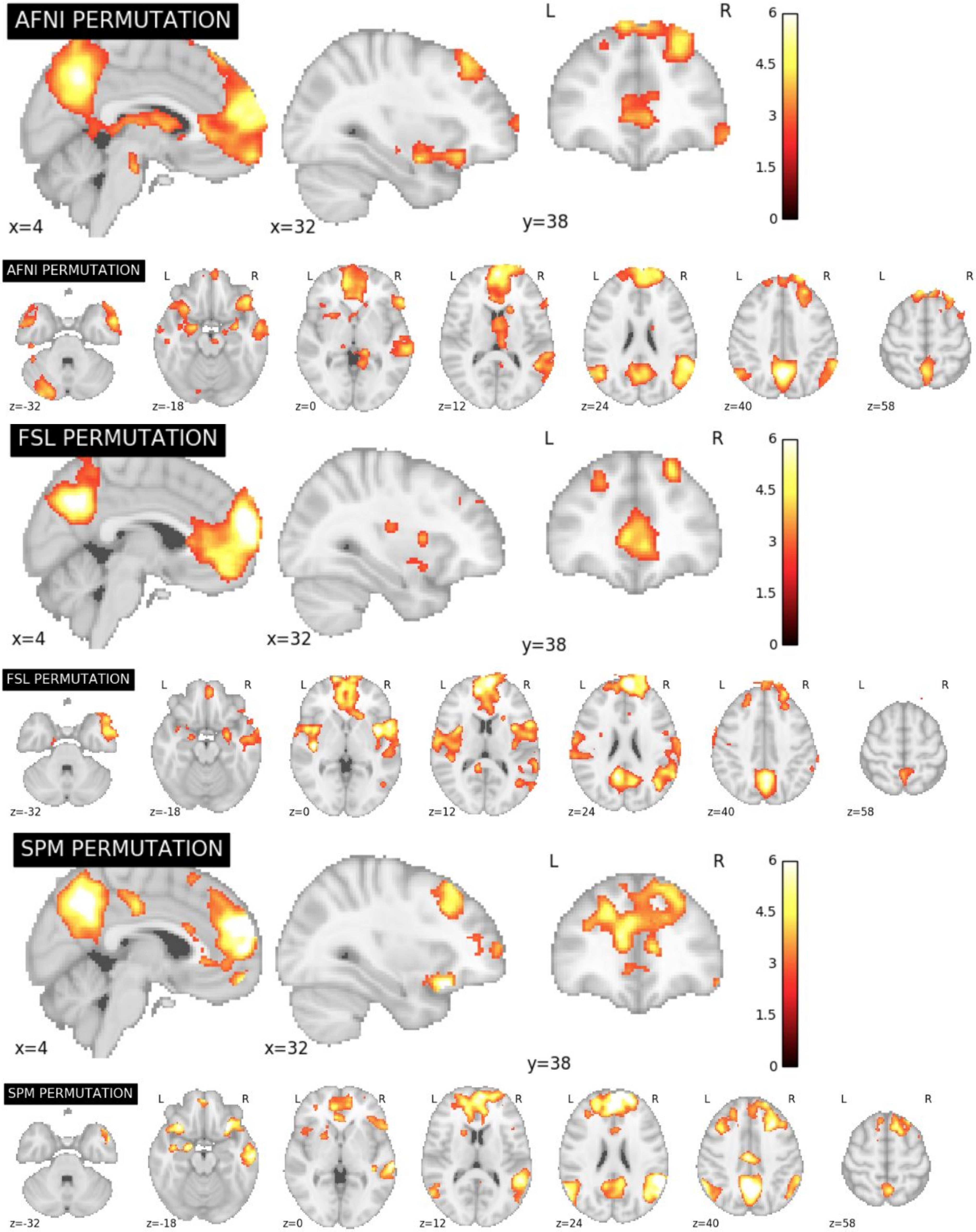

Figure S6. ds000120 Inter-Software Comparison, 5% FWE Clusterwise Inference

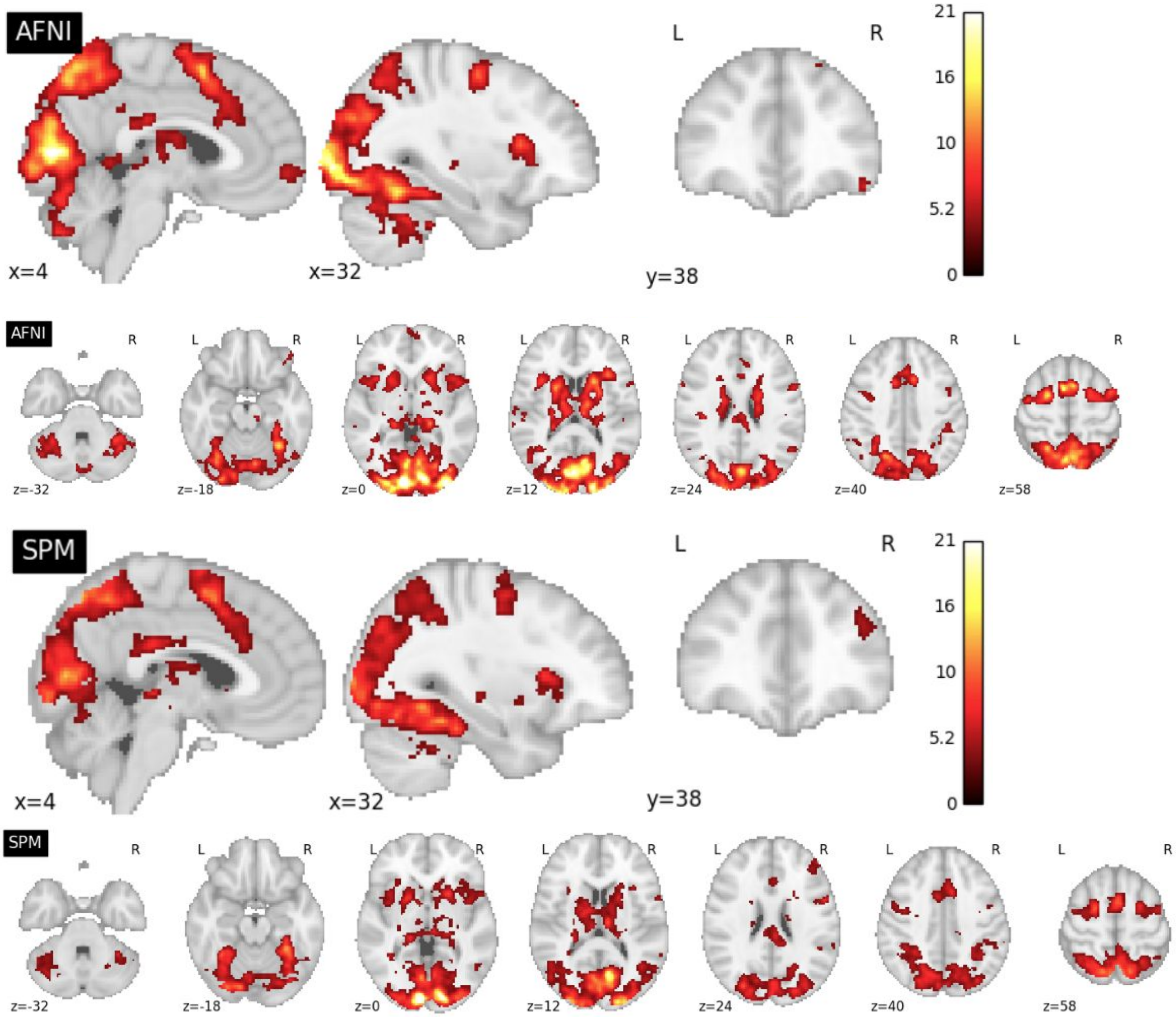

Figure S7. ds000001 Inter-Software Comparison, T-Statistic Maps

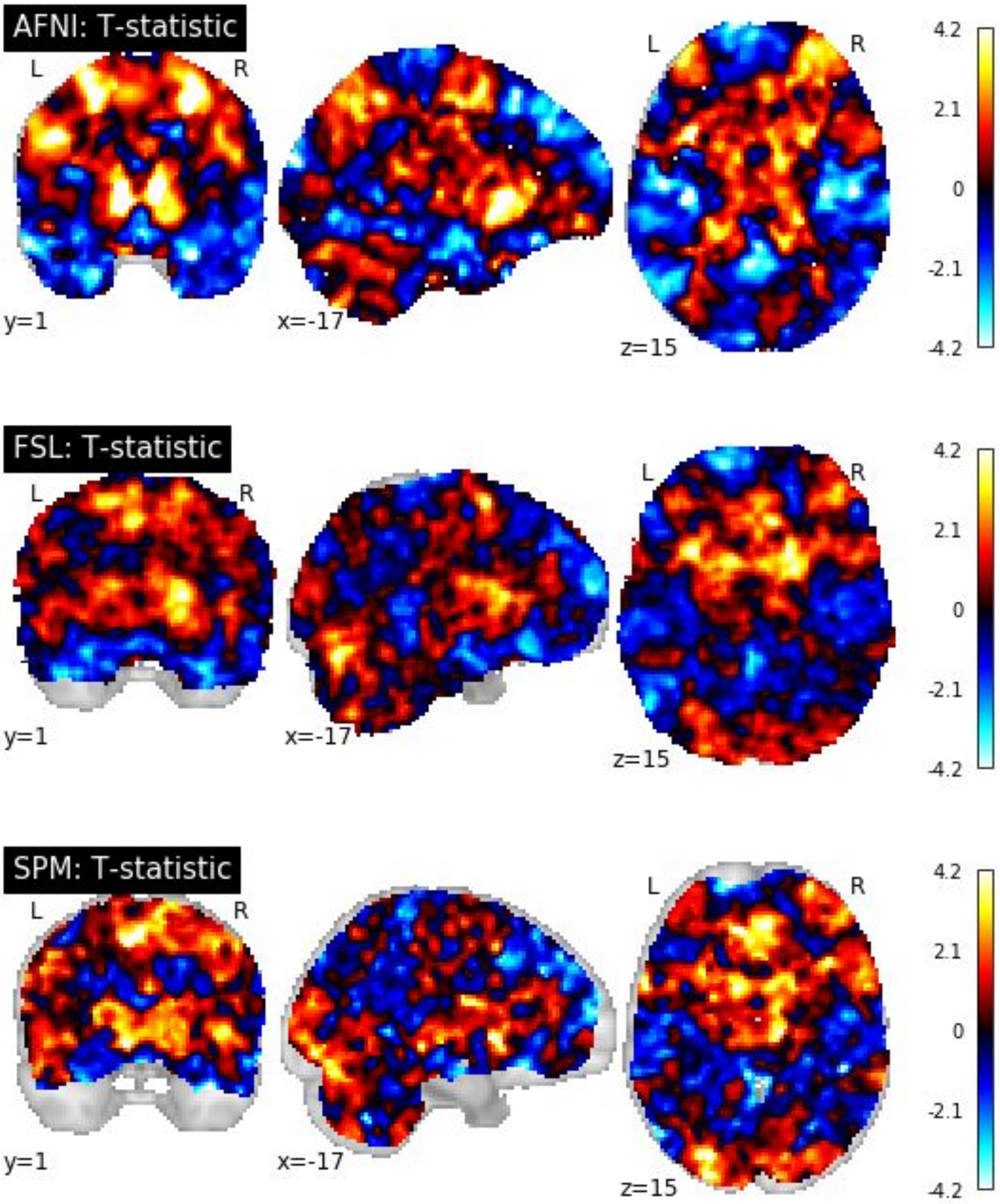

Figure S8. ds000001 Inter-Software Comparison, T-Statistic Maps from Permutation

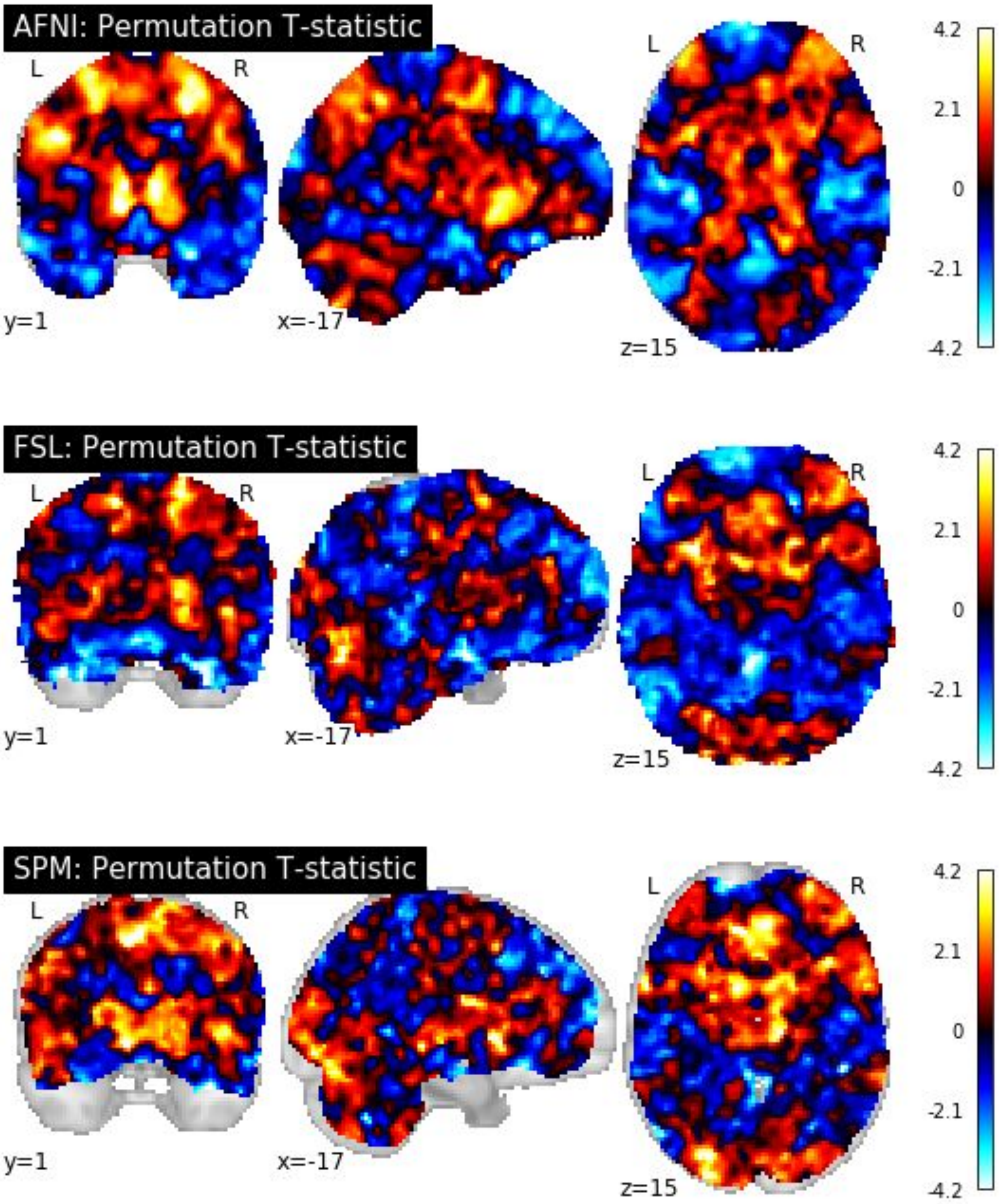

Figure S9. ds000109 Inter-Software Comparison, T-Statistic Maps

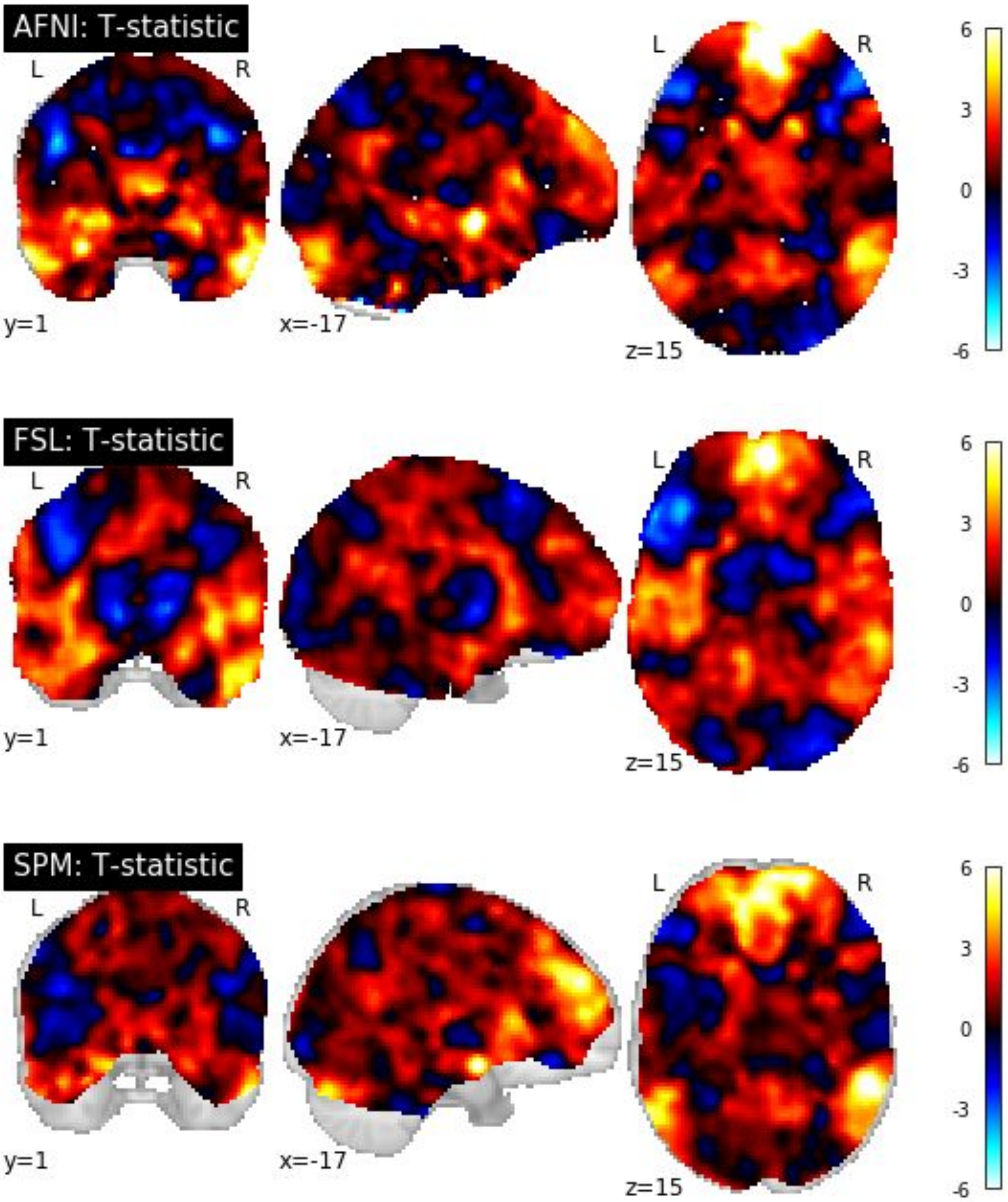

Figure S10. ds000109 Inter-Software Comparison, T-Statistic Maps from Permutation

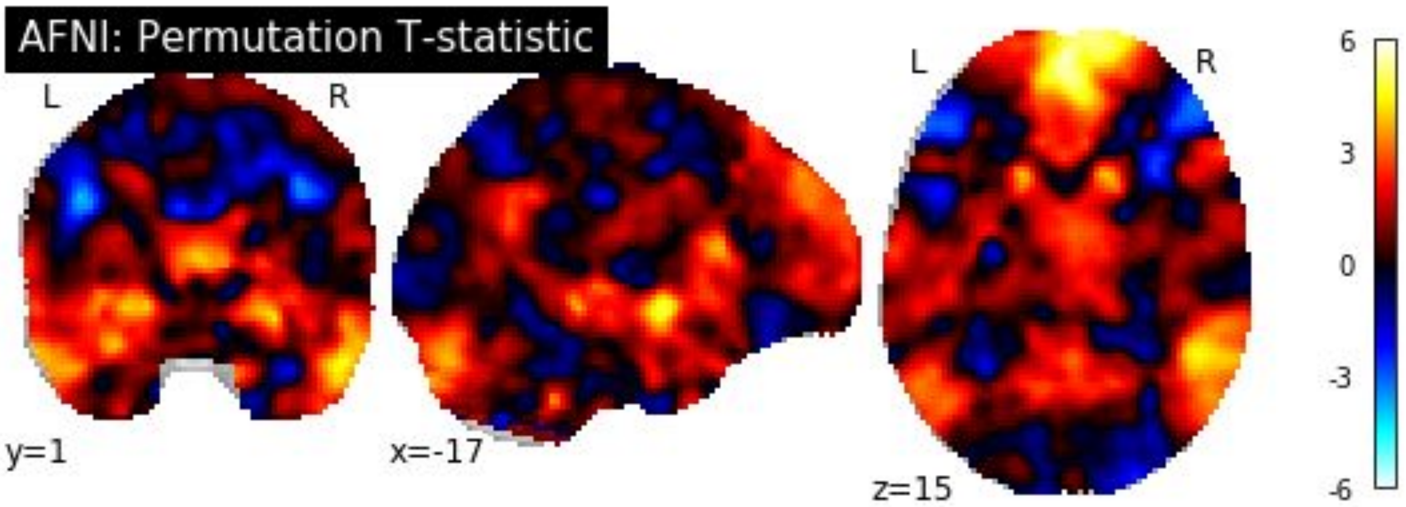


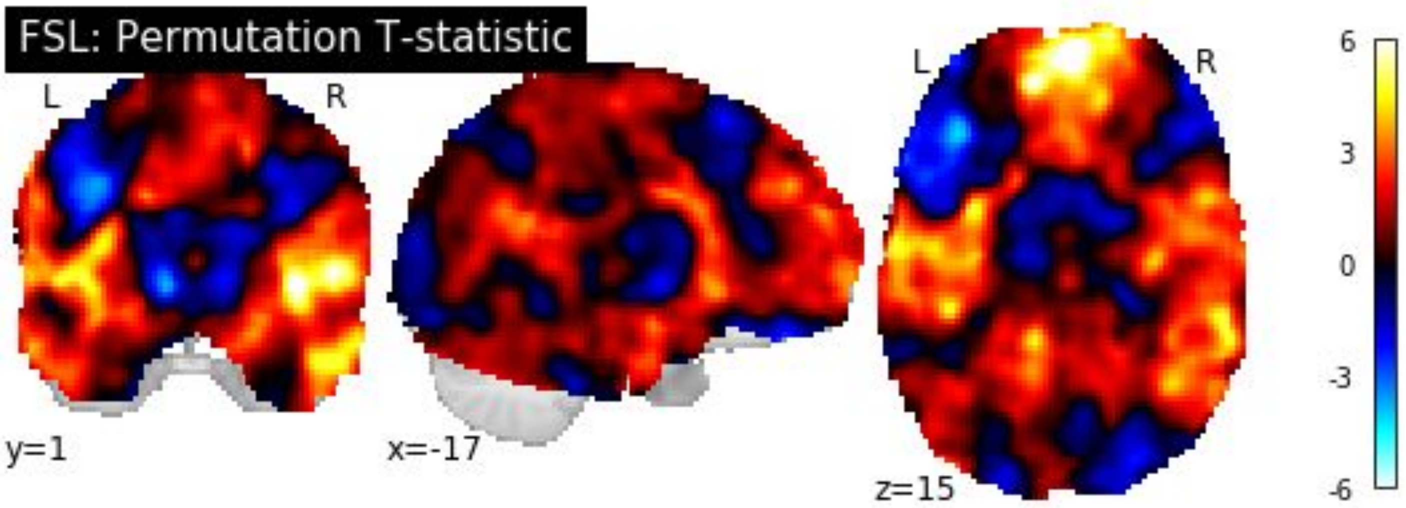


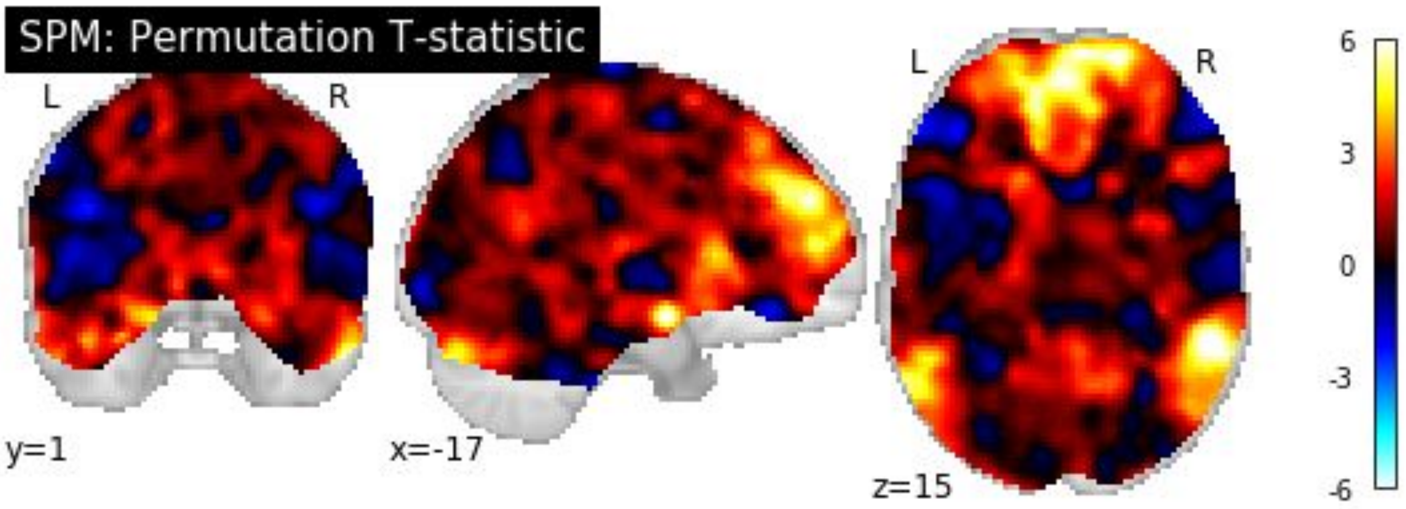

Figure S11. ds120 Inter-Software Comparison, F-Statistic Maps

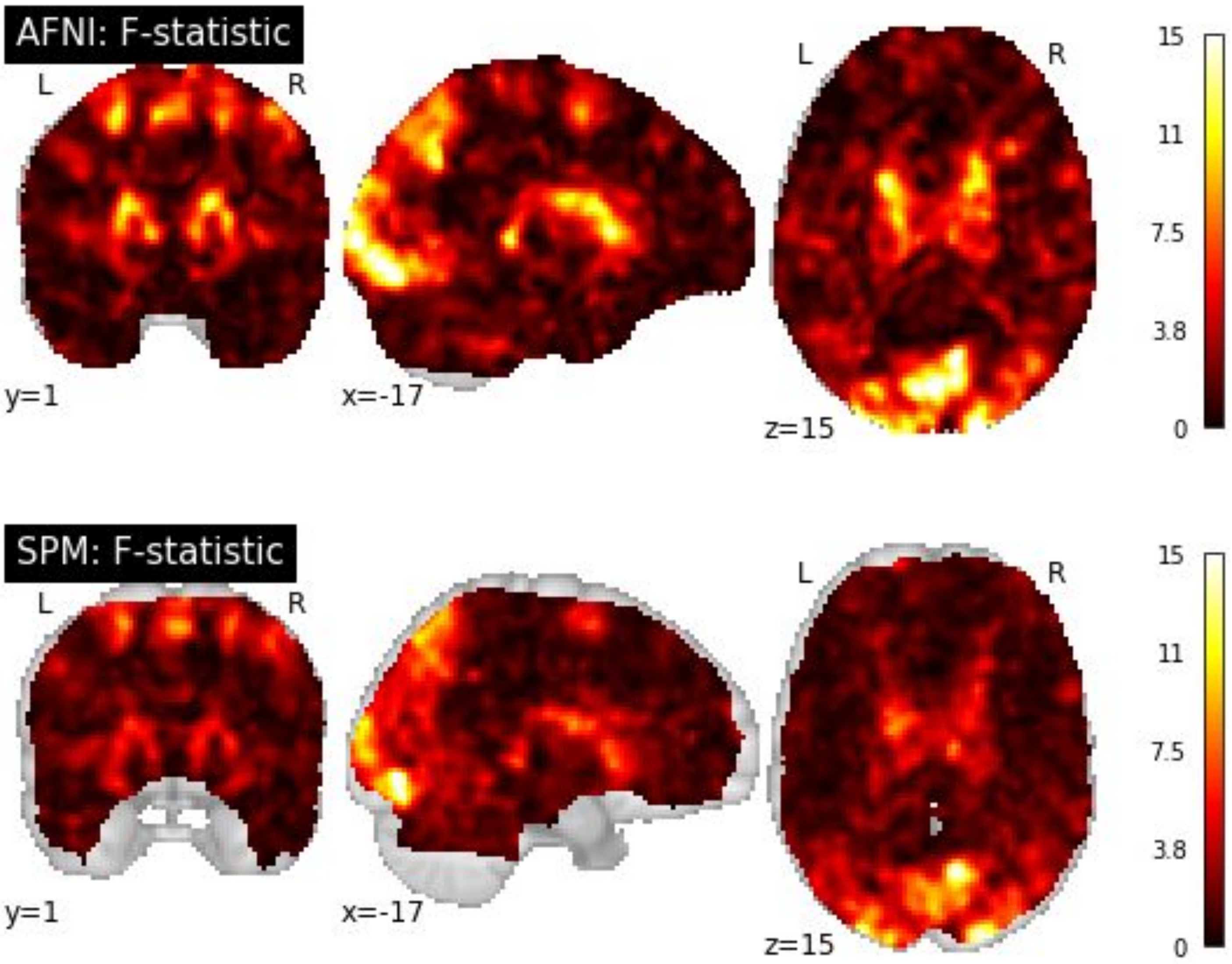

Figure S12. ds000120  Inter-Software Comparison, Euler Characteristic Curves for F-Statistic Maps

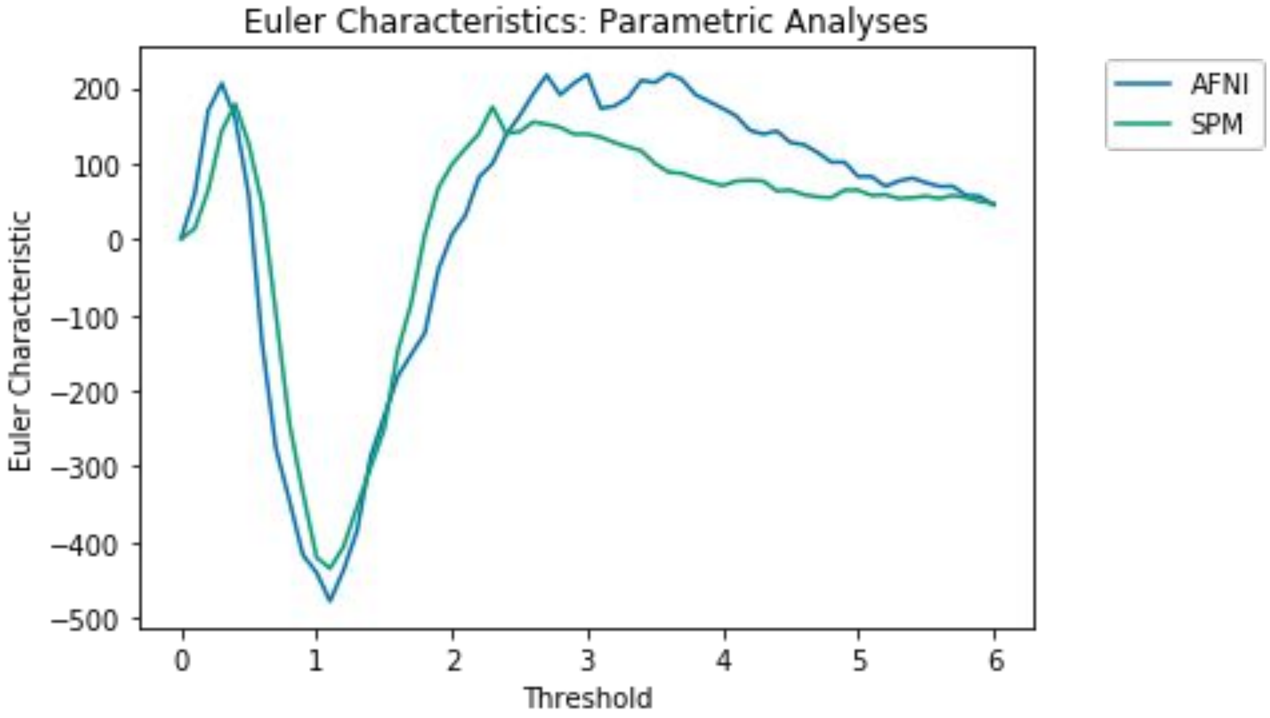
